# Toward simple, rapid, and deep plant proteome analysis with an in-cell proteomics strategy

**DOI:** 10.1101/2025.10.30.685699

**Authors:** Deji Adekanye, Meghana Kusuru, Jeffrey L Caplan, Yanbao Yu

## Abstract

Although modern mass spectrometry (MS) based proteomics has been broadly employed by the plant science community, proteomics analysis of plant tissues remains highly challenging to general plant biology laboratories, mainly due to the lack of effective and easily adaptable strategies that can process plant samples for MS analysis rapidly and reliably. Existing methods often involves lengthy mechanical disruption and protein extraction, which not only require large starting materials but also can cause significant variations, compromising quantitation accuracy. Here, we present an “in-cell proteomics” strategy and benchmark its performance against conventional processing methods using several model plants, including *Arabidopsis thaliana*, *Nicotiana benthamiana*, *Zea mays* and *Sorghum bicolor*. The comprehensive datasets from this study indicate that the on-filter in-cell (OFIC) processing-based “in-cell proteomics” strategy can eliminate cell lysis of plant leaves and pollen grains, and process proteins directly, enabling the identification of 9,000 to 12,000 proteins from leaves, and 7,000 to 9,000 proteins from pollen grains using one-shot LCMS. The OFIC strategy demonstrated broadly equivalent or better performance over conventional approaches in the context of simplicity, speed, digestion effectiveness, and quantitation accuracy. Applying the strategy to investigate plant leaf infection of *N. benthamiana* leaves revealed interesting signature proteins that respond differentially to host-adapted *Colletotrichum destructivum* and nonhost-adapted *Colletotrichum sublineola* fungus. The single-vessel approach bypasses cell disruption, substantially simplifying plant sample preparation while reducing the required sample input and enhancing proteomic sensitivity. It offered a generalizable approach to study plant leaf infection and other significant questions in plant biology and beyond.

## Introduction

Plants are fundamental for both food and human health, providing essential nutrients, supporting ecosystems, and contributing to overall well-being (Raskin et al., 2002; Martin et al., 2011). Liquid chromatography and mass spectrometry (LCMS) based proteomics have been employed broadly to study plant biology, playing a vital role in understanding plant protein functions, interactions, and responses to various stimuli in their environment (Mergner and Kuster, 2022; Geddes-McAlister and Uhrig, 2025). However, proteomic applications within plant sciences still remain under-represented. This is not only due to the high technical barrier to LCMS instrumentation, but also the lack of standardized, efficient, effective, and economical ways of processing plant samples for reliable and robust proteomics analysis. Unlike mammalian cells, a significant proportion of the plant cell is made up of a central vacuole surrounded by rigid cell walls, and these attributes pose substantial challenges to efficient protein extraction from plant cells, which is often the prerequisite for conventional bottom-up proteomics (Isaacson et al., 2006; Song et al., 2018). Another important attribute of plant cells for proteomics is the complexity and high dynamic range of the plant proteins, in addition to various interfering substances that include phenolic compounds, carbohydrates, pigments as well as a wide range of plant-specific and growth stage-dependent metabolites (Mergner and Kuster, 2022). Therefore, the protein extraction is a crucial step in plant proteomic analysis, which directly affects the reliability of downstream applications. Protein extraction in plants entails the disruption of the tough cell walls, followed by the removal of interfering substances using purification or protein precipitation techniques (Isaacson et al., 2006; Mikulášek et al., 2021). Additionally, the depletion of high-abundance proteins, such as RuBisCO, can be performed to enhance the detection of low-abundance proteins (Kim et al., 2013; Gupta et al., 2015). Disruption of plant cell walls is often achieved by mechanical disruption techniques like bead milling, chopping in a blender, or liquid nitrogen grinding with a mortar and pestle. Unfortunately, these procedures are not only time consuming but also can cause protein degradation, variable extraction efficiency, or significant protein loss under suboptimal conditions (Isaacson et al., 2006). Detergent-based extraction facilitates the solubilization of membrane proteins, improving recovery, but detergents interfere with LCMS analysis and necessitate additional purification steps, which are again not only lengthy but also can introduce technical variabilities, compromising accuracy for quantitative proteomics analysis (Wu et al., 2014).

On the other hand, to obtain high proteome coverage and to detect biologically relevant low-abundant proteins, deep sampling strategies are often applied, such as on-line or off-line two-dimensional LC. For instance, using strong cation exchange LC and hydrophilic strong anion exchange LC, a range of 18 to 30 fractions were measured separately by MS to identify 7,889, 11,690, and nearly 15,000 proteins from *Arabidopsis* leaves (Song et al., 2018; Zhang et al., 2019; Mergner et al., 2020). Other pre-fractionation techniques, such as long on-line LC gradients, high-field asymmetric waveform ion mobility spectrometry (FAIMS)-based gas phase fractionation, and sequential window MS1 acquisition, have been explored as well to reduce sample complexity while allowing single-shot analysis (Meier et al., 2018; Bian et al., 2021; Gallo et al., 2023). However, these alternatives may substantially increase the data acquisition and analysis time, reduce measurement reproducibility, and thus the quantitation accuracy. Therefore, there is an urgent unmet need to develop an effective but simple sample processing approach for plant proteomics.

We recently developed an on-filter in-cell (OFIC) processing-based E4technolgy, which eliminates cell lysis and protein extraction, by digesting proteins directly in the cells after a simple methanol-based fixation (Martin et al., 2024). Methanol treatment makes cell membranes permeable and kills cells by aggregating cellular proteins, allowing proteolytic enzymes to penetrate into the cells and cleave proteins (Hatano et al., 2023). We have validated the method previously using mammalian cells and *Xenopus* embryos (Sun et al., 2025a; Xu et al., 2025) and other types of hard-to-lysis samples, such as yeast cells (Martin et al., 2024) and whole *Caenorhabditis elegans* animals (Elsayyid et al., 2025). For instance, without grinding or homogenization of the *C. elegans*, we were able to digest proteins directly in the fixed worms, and identify over 8,000 proteins from as few as 50 worms. Single-worm proteomics became technically feasible as well. Such an “in-cell proteomics” approach integrates all the sample preparation steps into a single filter device, including reduction and alkylation, protein digestion, and peptide cleanup, thus drastically simplifying sample processing and minimizing sample loss. Compared to traditional lysate-based digestion methods, the OFIC approach does not compromise digestion efficiency while it enhances the proteomics sensitivity and quantitation accuracy. Encouraged by the successful experimental evidence, in this study, we attempt to test the “in-cell digestion” idea using plant tissues, which represent one of the most challenging samples for proteomics analysis. We utilized several different types of plants and tissue samples to verify the applicability of the method, including leaves from *Arabidopsis thaliana*, *Nicotiana benthamiana*, *Zea mays* and *Sorghum bicolor*, and pollen from *A. thaliana* and *N. benthamiana.* We further applied the method to investigate a biologically significant question, how does the proteome of *N. benthamiana* leaves dynamically change upon inoculation with host-adapted *Colletotrichum destructivum* and nonhost-adapted *Colletotrichum sublineola* fungus. With in-cell proteomics, we were able to gain insights into how adapted *C. destructivum* suppresses or induces different *N. benthamiana* defense responses.

## Results

### Benchmarking in-cell proteomics analysis of sorghum leaf

We first set out to examine whether methanol can render plant leaf tissues cleavable by proteolytic enzymes. Brightfield imaging data suggested that, upon methanol fixation, sorghum leaf sections showed a progressive clearing of chlorophyll over 30 minutes (Supplementary Figure 1A). To examine the cellular details, we labeled the cell wall with calcofluor white (CW) dye and conducted multiphoton microscopy (Supplementary Figure 1B). Methanol clearing resulted in improved, deeper staining and imaging of the cell wall of sorghum leaf sections. The tissue appeared to be more fragile after methanol treatment, and surface damages from handling the sample during the imaging were observed. Interestingly, cracks in the cell were also observed at the junctions of sorghum cells (Supplementary Figure 1B). The changes in the cell wall may make the cell wall more porous and penetrable for proteolytic enzymes, since we recently showed that methanol fixation facilitated penetration of trypsin through the thick cuticles of *C. elegans*, and enabled direct digestion of proteins inside the intact worms (Elsayyid et al., 2025). Next, we examined the effect of exchanging and vortexing with fresh methanol every five minutes for 30 minutes (six exchanges) (Supplementary Figure 2A). The multiple methanol exchanges depleted the green chlorophyll completely, thus greatly reduced the potential interferences with downstream LCMS acquisition by residual chlorophyll and primary and secondary metabolites in leaf tissue (Mergner and Kuster, 2022), and improve the identification of low-abundance proteins, as will be discussed below.

Physical disruption, such as sonication, bead beating, grinding or blending, followed by protein extraction with SDS buffer, has been commonly used for plant leaf proteomics (Wang et al., 2018; Mikulášek et al., 2021; Balotf et al., 2022). On the other hand, very recently TFA was shown to be able to dissolve hard-to-lyse cells such as gram-positive bacteria and microbiome samples (Doellinger et al., 2020). To our knowledge, TFA has not yet been applied to explore plant leaf proteomics. In this study, we thoroughly compared OFIC processing with SDS- and TFA-lysate based digestion methods. We took five proximal leaves derived from 4-5-week-old sorghum plants, and sliced them into approximately 1-2 cm^2^ sections. We treated each section as one biological replicate, and further cut it into 1-2 mm small pieces (Supplementary Figure 2B), which were then transferred to E3filters followed by methanol fixation, reduction and alkylation, protein digestion, and single-shot LCMS analysis (**Supplementary Protocol**). The SDS- and TFA-based digestion experiments were performed in parallel (**Figure 1**). Our data indicated that after a simple methanol fixation, surprisingly, the leaf slices were readily available for digestion, as evidenced by over 10,000 protein groups and nearly 90,000 peptide identifications from as little as ¼ of a single leaf tissue (**Figure 2, A-B; Supplementary Table S1**). Only less than 1% of the proteins were exclusively identified by the OFIC method; the vast majority of them were also identified by at least one of the lysate-based digestion methods (Supplementary Figure 3A), suggesting that in-cell processing is not prone to proteome-wide bias or under-representation. Because the OFIC method drastically streamlined proteomics sample preparation, we observed minimal sample requirement, higher reproducibility and quantitative accuracy compared to the lysate-based processing methods, which is consistent with what we observed from the worm experiments (Elsayyid et al., 2025). The coefficient of variance (CV) of protein groups was around 15% and the Pearson’s correlation was near 0.97 between replicate experiments, both of which were higher than SDS- and TFA-lysate-based methods (**Figure 2C; Supplementary** Figure 3B). Even though the proteins remained inside the mostly intact leaf tissue, the enzymatic digestion was minimally affected. Less than 18% of the total number of detected peptides carried miscleaved sites, and their abundance was low, accounting for only 6-7% of the total peptide intensity (**Figure 2D; Supplementary** Figure 3C).

**Figure 1.**
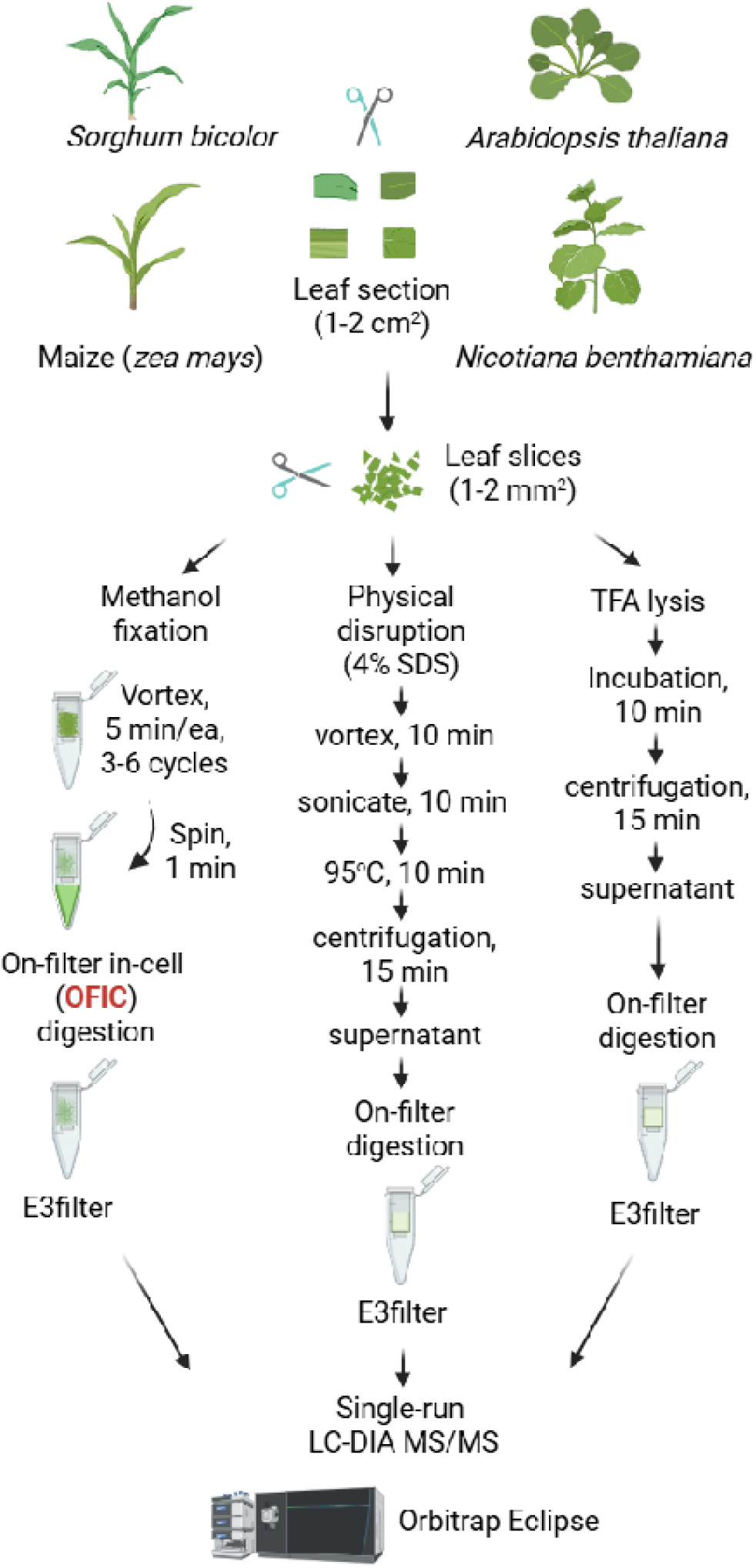
Illustration of plant proteomics workflows investigated in this study. Plant leaf tissues were first sliced into small sections, and then were subjected to methanol-based chlorophyll depletion and fixation, and on-filter in-cell digestion. In parallel, leaf slices were lysed with SDS-based buffer and strong acid TFA followed by on-filter digestion. A single-shot LCMS analysis was performed using Orbitrap Eclipse with FAIMS interface.

**Figure 2.**
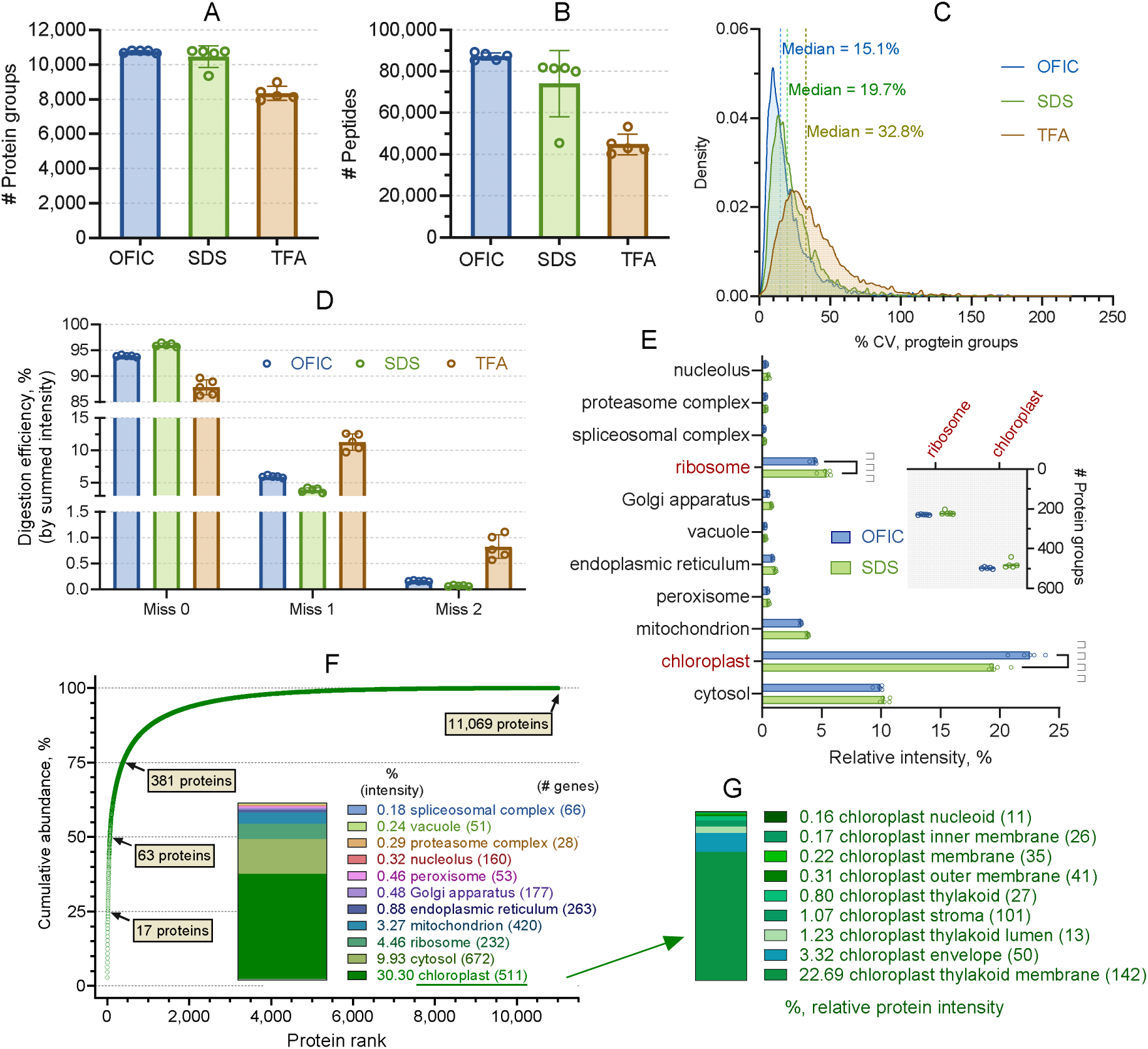
Benchmarking in-cell proteomics analysis of sorghum leaf. (**A-B**) Number of protein groups and peptides identified by the three processing methods, respectively. Error bars represent five biological replicates. (**C**) Profiles of protein groups coefficient of variation. (**D**) Digestion efficiency assessment based on peptide intensity. (**E**) Cellular compartment analysis of the proteins derived from OFIC and SDS-based processing method. (**F**) Cumulative protein abundance analysis of sorghum proteome derived from OFIC processing. The numbers of proteins account for 25%, 50%, and 75% of total protein mass are indicated in the plot. Inner diagram shows relative abundance of the eleven protein categories. (**G**) Histogram shows the relative abundance of chloroplast-associated proteins. The number of proteins for each category was indicated in the brackets.

We next assessed the dynamic range of the sorghum leaf proteome derived from OFIC digestion. Taking the median of intensity-based absolute quantification (iBAQ) values of the five replicates, the OFIC approach identified a total of 11,069 sorghum proteins, which spanned nearly seven orders of magnitude, and 63 proteins alone accounted for nearly half of the total protein mass (**Figure 2F; Supplementary** Figure 3D). Due to the unsatisfactory performance of TFA lysis, we excluded the TFA-derived proteome from further data analysis. In terms of the overall protein ontologies, both OFIC and SDS methods identified a similar quantity of proteins across a wide range of cellular compartments, such as cytosol, mitochondrion, and nucleolus (**Figure 2E**). Proteins related to ribosome and chloroplast showed quantitative variations, although no qualitative differences were seen between the two digestion methods (**Figure 2E, inner panel**). Rather than RuBisCo, the most abundant proteins in sorghum leaf appeared to be ATP synthase subunit alpha (aptA) followed by Germin-like protein (GLP), Plastocyanin, Cytochrome b559 subunit alpha (psbE), and Photosystem II D2 protein (psbD). As a C4 plant, this data makes sense because sorghum is known for its high photosynthetic efficiency and requires less Rubisco enzymes compared to C3 plants such as Arabidopsis (Sage, 2002). Interestingly, these top 5 most abundant proteins were all annotated as chloroplast localized proteins. The chloroplast is a hallmark organelle in plant cells, playing fundamental roles in photosynthesis, amino acid synthesis, lipid metabolism, and immune responses (Wang et al., 2023b). Our study identified over 500 chloroplastic proteins, which accounted for nearly 1/3 of the total protein mass (**Figure 2F, inner panel**). This data suggests the great dominance of chloroplast proteins in plants, which is not surprising. However, the results may still be underrepresented, as only 70% of sorghum proteins are annotated in the context of cellular compartment (CC) in the DAVID Knowledgebase. In contrast, over 99% of Arabidopsis proteins have clear compartmentation annotation in DAVID, and over 3,000 chloroplast associated proteins were identified in this study (as will be discussed later). Lastly, we examined the subclasses of the chloroplast proteins, in particular the chloroplast membrane proteins. In total, the OFIC digestion data mapped 142 thylakoid membrane proteins as well as 67 chloroplast inner and outer membranes, representing the most abundant category of proteins in sorghum leaf proteome (22.69%; **Figure 2G**). This data corroborates the subcellular localization distribution of the global leaf proteome and underscores the effectiveness of the OFIC method for plant membrane protein identification, despite the rigid protection of the cell wall and potential interference from the massive cellulosic matrix.

### In-cell proteomics analysis of maize, *Nicotiana benthamiana*, and Arabidopsis leaves

To investigate whether the OFIC method has any sample type restrictions or specificities, we further challenged the in-cell proteomics method with several other plants, including *Arabidopsis thaliana* (hereafter Arabidopsis), *zea mays* (hereafter maize), and *Nicotiana benthamiana*. We compared OFIC with the SDS lysate-based method whenever possible. At least three biological replicates were prepared for each method. To process leaf samples, we adapted the same procedures as illustrated in **Figure 1**. As expected, using a simple in-cell digestion method and a single-shot DIA MS/MS, we were able to identify approximately 9,600 proteins from Arabidopsis, 8,600 proteins maize, and 12,000 proteins from *N. benthamiana* (**Figure 3A-F; Supplementary Table S2, S3, and S4**). In line with our previous findings, the OFIC digestion outperformed SDS lysate-based digestion methods again in the context of protein and peptide hits, and is highly reproducible (**Figure 3G**) as well, ensuring excellent intensity correlation and quantitation accuracy.

**Figure 3.**
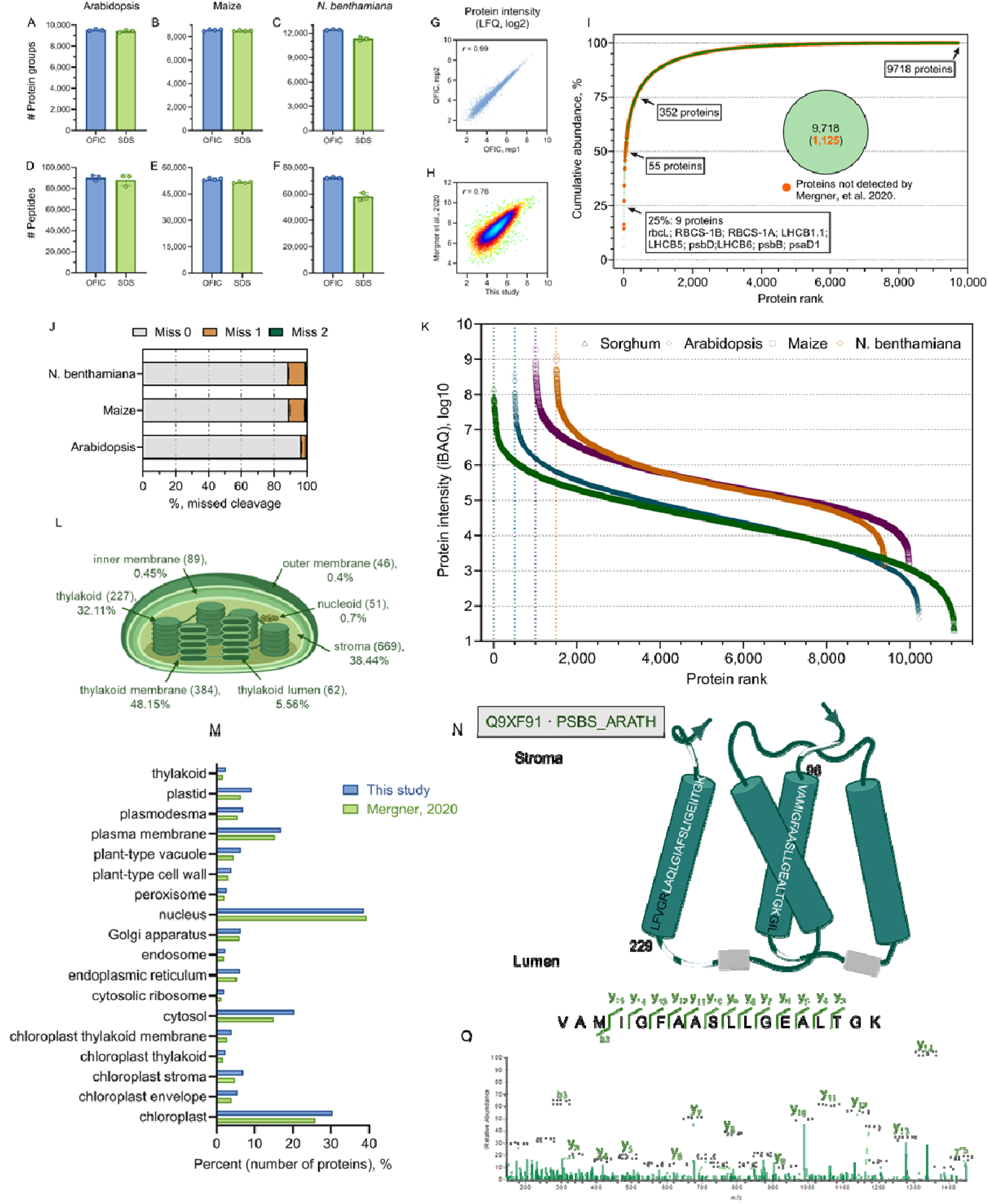
Applying OFIC digestion to proteomics analysis of other plant species. (**A-C**) Protein identifications from leaves of Arabidopsis, Maize, and *N. benthamiana*. (**D-F**) Peptide identifications of the three types of leaves. (**G**) Reproducibility assessment. Two replicates of OFIC digestion of Arabidopsis leaf samples were taken as a representative here. Pearson correlation r value is shown in the plot. (**H**) Density plot of the Arabidopsis proteins obtained from this study (OFIC digestion) and the study by Mergner *et al*. Protein group quantity (log2-transformed) was used for the plot. Pearson correlation r value is shown in the plot. (**I**) Cumulative abundance plot of Arabidopsis proteome obtained by OFIC digestion. The top 9 proteins in the first quarter (25%) as well as the number of proteins in the other quarters were indicated in the plot. The pie shows the total number of proteins from OFIC method, and the number of proteins that were exclusively identified by OFIC. (**J**) Digestion efficiency assessment of OFIC processing for the three types of leaves. (**K**) Dynamic range plot of the proteomes derived from OFIC processing of the four types of leaves. Inner panel shows the top 10 most abundant proteins of each leaf. (**L**) Quantitative proteomics of Arabidopsis chloroplast. Chloroplast proteins and their relative abundance are shown. Numbers in the brackets indicate the number of proteins of each category. (**M**) Cellular compartment enrichment analysis of the Arabidopsis proteome obtained from this study and the study by Mergner et al. (**N**) Transmembrane protein identification by OFIC method. Protein PSBS is shown as a representative. The peptides that were identified by OFIC are shown as while lines (outside of TM domain) and amino acid sequences (inside TM domain), respectively. A representative MS/MS spectrum is included as panel (**O**).

Maize leaf proteomics has been studied quite extensively. Over 12,000 proteins have been reported using 2D or 3D LC fractionation and 20-40 LCMS injections (Gao et al., 2008; Facette et al., 2013). *N. benthamiana* leaf proteomics applications appeared to be relatively limited regarding the depth, covering only less than 6,000 proteins with LC or gel-based pre-fractionation (Shen et al., 2022; Zheng et al., 2024). Arabidopsis is the foremost model plant species with a well-annotated genome. Mergner and coworkers reported the identification of approximately 15,000 Arabidopsis leaf proteins using exhaustive sampling strategies (i.e., multi-shot LCMS with 24 fractions) (Mergner and Kuster, 2022). With single-shot LCMS, Mikulášek et al. reported around 5,000 protein hits from SDS lysate of whole Arabidopsis leaf (Mikulášek et al., 2021). In this study, we utilized a much simpler and more rapid approach but achieved equivalent or deeper proteome coverage. Regarding the effectiveness of in-cell digestion, we observed generally good to excellent efficiency for all three types of leaves (**Figure 3J**), which is consistent with digestion data from sorghum leaf described above and *C. elegans* worms reported recently. Meanwhile, massive overlaps can be seen between the proteins obtained from in-cell digestion and the proteins derived from lysate-based processing, either in this study or from other studies (Supplementary Figure 4**, A-D**). Quantitatively, both the maize and the *N. benthamiana* proteome span over six orders of magnitude, suggesting broadly deep coverage of these two hard-to-lysis leaf tissues by the in-cell proteomics approach. On the other hand, comparing the Arabidopsis data obtained from this study and Mergner’s (Mergner and Kuster, 2022), we have also observed generally good correlation and comparable distribution of subcellular localizations with minor variations from cytosolic, chloroplastic, and mitochondrial proteins (**Figure 3, H** and **M**). In summary, these data suggested minimal quantitative bias of the in-cell proteomics approach and demonstrated the broad applicability of the in-cell digestion method for proteomic analysis of plant leaves.

We then took a further look at the OFIC-derived Arabidopsis leaf proteome, which spanned nearly seven orders of magnitude, and agrees well with previous reports (Mergner and Kuster, 2022). According to DAVID annotation, approximately 3,050 proteins were classified as chloroplastic proteins, which was the second largest category (slightly less than the 3,800 nuclear proteins) and accounted for over 78% of total Arabidopsis leaf protein mass. Interestingly, 384 chloroplast thylakoid membrane proteins, as classified by DAVID database, contributed to nearly half (48.2%) of the overall leaf protein mass (**Figure 3L**). This data underlined the potential of in-cell digestion strategy for deep chloroplast proteomics analysis. Here, we highlighted one protein to showcase that the “in-cell proteomics” approach can not only identify transmembrane proteins but also their transmembrane domains. Photosystem II 22 kDa protein (PSBS) is a multi-pass chloroplast thylakoid membrane protein. With in-cell digestion, seven peptides were identified for this protein, including two sequences in the stroma region, three in the lumen region, and another two in the transmembrane regions (TM1 and TM4; **Figure 3, N-O**). There was clear evidence that the cleavages occurred at the embedded sites in the lipid bilayer. These data support our hypothesis that methanol fixation may solubilize phospholipids and facilitate the tryptic cleavage, thereby enabling successful identification of integral membrane proteins. From a quantitation standpoint, the three most abundant proteins in Arabidopsis leaves are RuBisCO enzymes (RbcL, RbcS-1B, and RbcS-1A) that contributed to over 14% of the total protein mass (**Figure 3I**), which is expected for a C3 plant such as Arabidopsis. The overwhelmingly high abundance of RuBisCo proteins has been a major challenge for plant leaf proteomics, and a depletion strategy has been developed to identify low-abundance proteins (Fröhlich et al., 2012; Gupta and Kim, 2015; Tola and Missihoun, 2023). Here, our data showed that even without RuBisCO depletion, the “in-cell proteomics” strategy in combination with DIA LCMS is capable of deep proteome analysis of leaves.

### Applying in-cell proteomics to plant pollen

We next asked if the in-cell approach would be used to digest non-leaf samples. We took pollen grains as a proof-of-principle sample to demonstrate the versatility and power of the approach. For the pollen sample preparation, rather than E3filter, we utilized a smaller but functional device, E4tip (**Figure 4A; Supplementary Protocol**), which is embedded with reversed phase-based chromatographic resins and is capable of peptide binding. We recently showed that this single integrated tip can process low-number intact cells and *C. elegans* worms for proteomics analysis (Martin et al., 2024; Elsayyid et al., 2025). Previously, pollen proteomics studies required that the grains to be ground with a chilled mortar and pestle or disrupted with bead beater, to break the rigid pollen coat and extract proteins (Dai et al., 2007; Rejón et al., 2016; Darnhofer et al., 2021). The procedures tend to be lengthy, involve significant sample loss, and often require large inputs. Here, we reason that pollen grains are microns-to-millimeters in particle size and similar in shape to large-sized cells, thus may be processed in E4tips. For each pollen digestion experiment, we used around 1/10 the pollen grains collected from a single *N. benthamiana* flower, and grains pooled from two Arabidopsis flowers. Based on sample availability, we performed three replicate digestion experiments with *N. benthamiana* pollen, and one for Arabidopsis pollen. Surprisingly again, the methanol fixation enabled direct digestion of proteins in the intact pollen grains, leading to approximately 8,500 and 6,400 protein identifications from *N. benthamiana* and Arabidopsis pollen, respectively (**Figure 4, B-C; Supplementary Table S5 and S6**). The digestion efficiency was relatively lower in the pollen samples compared to the leaf samples, but still greater than 80% of the peptides were completely digested (**Figure 4D**), outperforming some of the commonly used filter-based digestion approaches (Ludwig et al., 2018).

**Figure 4.**
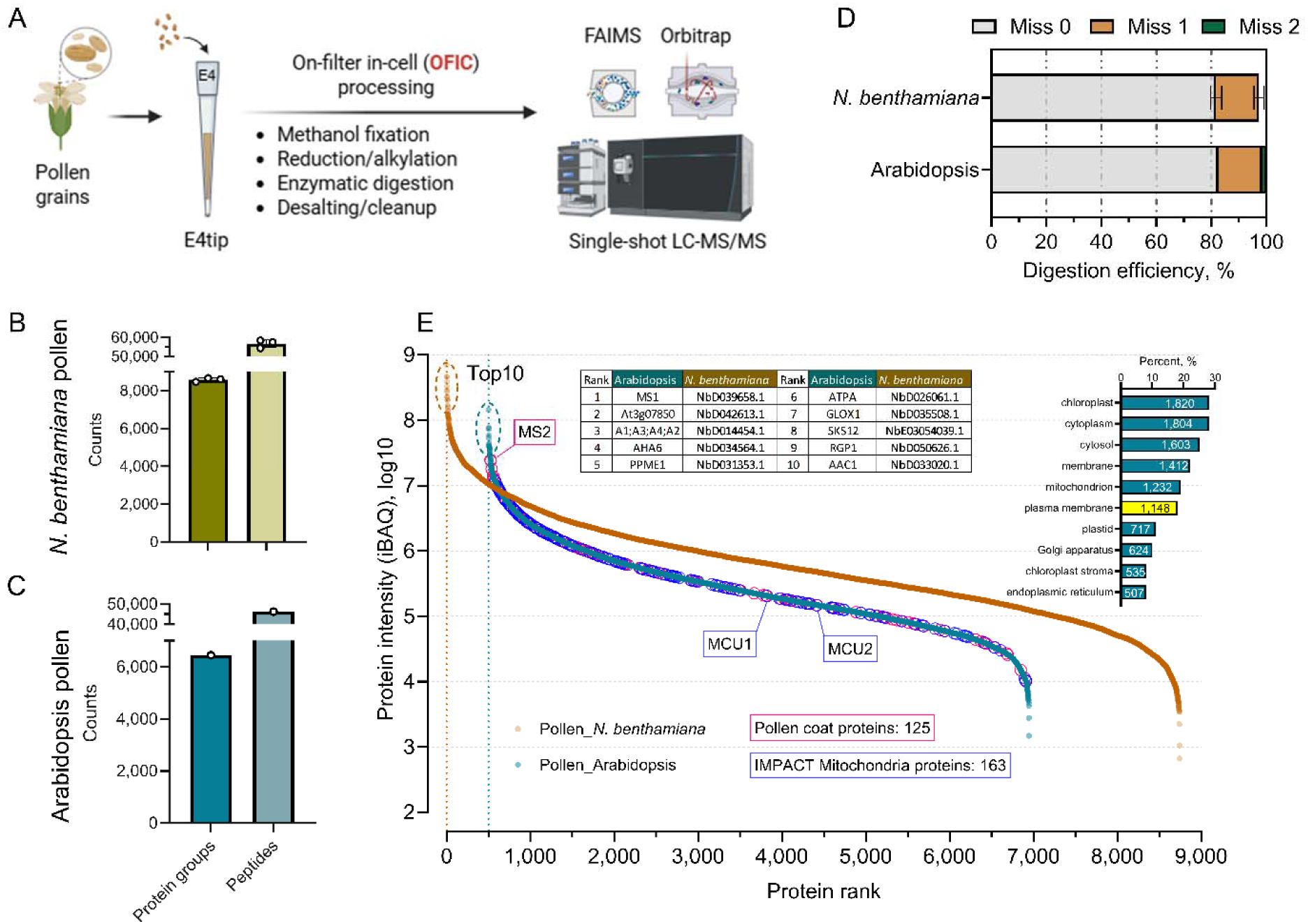
In-cell proteomics analysis of plant pollen. (**A**) Illustrative workflow. (**B-C**) Protein and peptide identifications from Nicotiana pollen and Arabidopsis pollen, respectively. (**D**) Digestion efficiency assessment. (**E**) Protein ranking plot. Inner table shows the top 10 most abundant proteins in the two plant pollen proteomes. The inner bar diagram shows the top 10 most enriched cellular compartment categories.

Pollen methionine synthase 1 (MS1) is a plant enzyme that synthesizes methionine for pollen development, particularly for the activation of calcium channels important for seed germination and pollen tube growth. Interestingly, MS1 protein was the most abundant protein in both Arabidopsis and *N. benthamiana* pollens we investigated (**Figure 4E**). Noteworthy is that many (∼42%) of the pollen coat proteins reported in a recent study were also identified by the OFIC approach (Wang et al., 2023a), such as MS2, an essential component of the pollen wall involved in the synthesis of fatty alcohols. Pollen development is extremely energy-demanding, and mitochondrial proteins are thought to play crucial roles in this process. A recent proteomics study of enriched mitochondria using IMPACT strategy identified approximately 200 proteins enriched in pollen mitochondria (Boussardon et al., 2025), and the OFIC approach detected 80% of them, highlighted by mitochondrial calcium uptake 1 and 2 proteins (MCU1, MCU2). A broader GO enrichment analysis suggested that nearly 1,148 pollen proteins (∼18%) were annotated as plasma membrane proteins. The above data underscored again the power of the in-cell proteomics approach, which can cleave proteins effectively in intact pollen grains.

### Proteomic investigation of *N. benthamiana* leaves inoculated with *Colletotrichum*

Having established a robust “in-cell proteomics” workflow for rapid and deep proteome analysis of plant tissues, we next sought to provide proof-of-principle evidence of this workflow for answering critical biological questions. For this purpose, we inoculated the *N. benthamiana* leaves with two strains of fungus that exhibit different host specificity, one was the nonhost-adapted *C. sublineola* that can invoke nonhost resistance (NHR), and the other was the host-adapted *C. destructivum* that infects and causes anthracnose disease. We processed them for proteomics analysis using the in-cell approach described above (**Figure 5A; Supplementary Table S7**). Experimentally, we identified on average 9,300 proteins from each one of the three replicates per group, totaling approximately 9,500 proteins with minimal missing values and high reproducibility (**Figure 5B; Supplementary** Figure 5A-B). Interestingly, although the leaf proteomes displayed largely similar patterns upon infection, their global profiles were already distinguishable according to a straightforward principal component analysis (Supplementary Figure 5C), suggesting that the single-run, in-cell proteomics approach was able to capture proteome-wide subtle alterations upon fungal inoculation.

**Figure 5.**
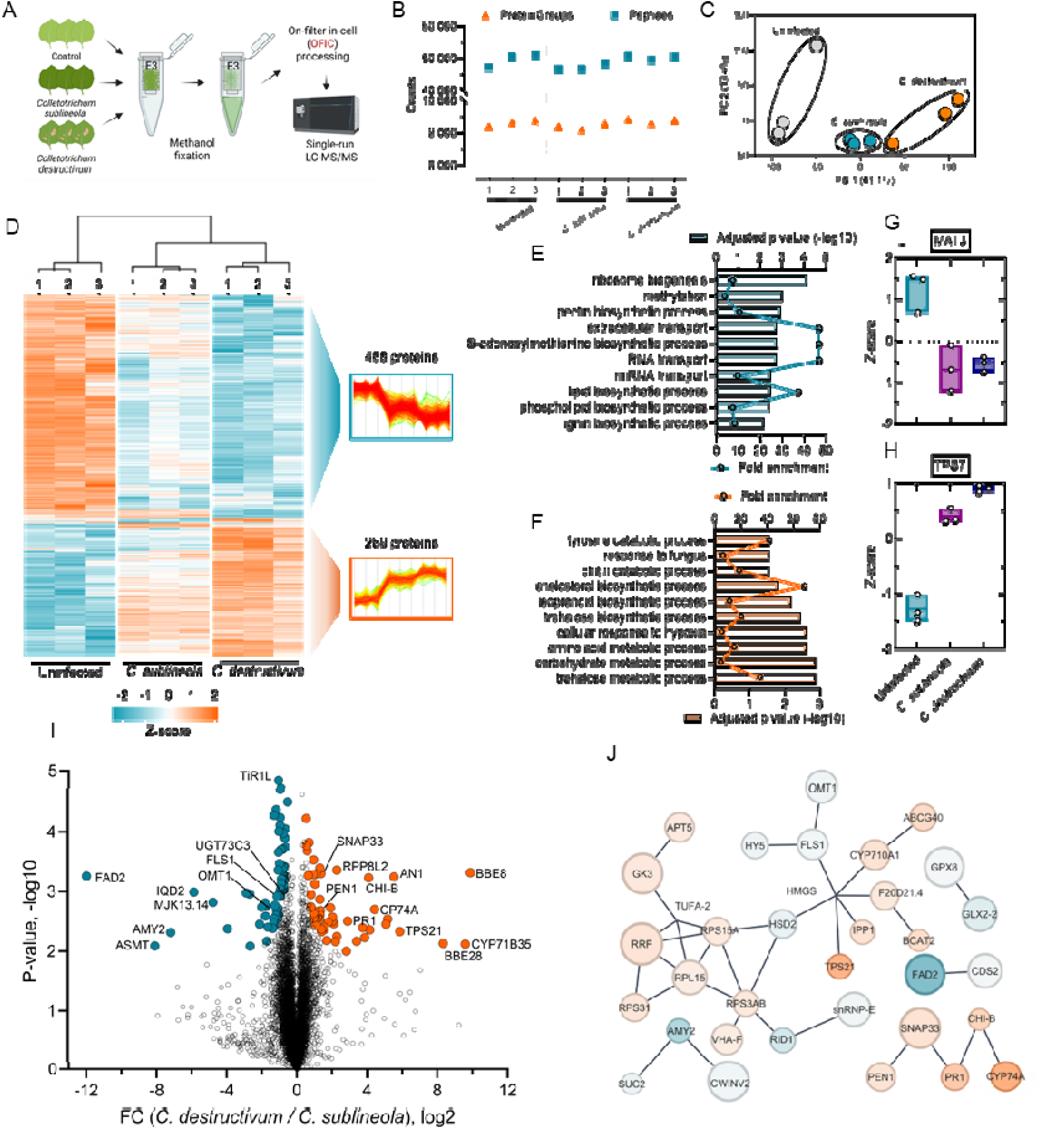
Applying in-cell proteomics to examine plant infection and immune response. (**A**) Experimental flowchart. (**B**) Protein and peptide identifications of the three groups. Three biological replicates were included for each group. (**C**) Principal component analysis. (**D**) Heatmap of ANOVA significant proteins (Permutation FDR 0.01, S0 = 0.5) among the three conditions. The proteins have been filtered out of protein identified by single peptides. (**E-F**) Gene Ontology enrichment analysis of the two clusters highlighted in the heatmap. (**G-H**) Two representative proteins that showed opposite patterns upon infection. (**I**) Volcano plot. The pairwise t-test (FDR 0.05, S0 = 0.1) of the two infected groups. (**J**) Protein network analysis. The differentially expressed proteins that showed significance in panel I were used to build the STRING network in Cytoscape.

We looked further into the quantitation data to learn more about the molecular defense mechanisms and metabolic alterations in response to fungus infection. A multi-sample ANOVA test (p < 0.05) led to ∼2,800 significantly differentiated proteins among the three groups, indicating a system-level proteome remodeling triggered by the infection (Supplementary Figure 5C). We then narrowed down the candidates utilizing a more stringent cutoff (Permutation FDR 0.01, stabilizing constant S0=0.5) for comparison, and also required the candidates to be matched by at least two unique peptides. The analysis resulted in 819 significant proteins among the three conditions, which were grouped in two major clusters, with 488 proteins in one cluster showed broadly declined expression upon fungal inoculation, and 289 proteins in the other showed up-regulation (**Figure 5D**). Multiple metabolic processes were found to be up-regulated upon fungus infection. Among them, the trehalose metabolic process showed the highest significance (with high fold enrichment). Trehalose is a sugar that supports fungal pathogens’ infection of plants by promoting appressorium development and maintaining cellular turgor, which are crucial for penetrating plant tissue (Foster et al., 2003). Ten trehalose-phosphate synthases were identified from this experiment, and four of them showed a significant increase, including TSP10, TSP11, TSP6, and TSP7 (**Figure 5F**). The down-regulated proteins are mainly involved in ribosome biogenesis, methylation, and pectin biosynthetic process. While methylation can influence ribosome biogenesis by altering the expression of key nucleolus proteins, their declined expression may imply a reduced level of protein translation or synthesis. This finding was corroborated with the decline of the S-adenosylmethionine (SAM) biosynthetic process. The adenosyltransferase (MAT) proteins play crucial roles in one-carbon metabolism, providing the methyl group for transmethylation reactions (Hanson and Roje, 2001). At least three MAT proteins were mapped in our study, including MAT3 (or SAMS3; **Figure 5E**), which has been reported previously to be targeted and manipulated by certain plant pathogens to weaken the host’s defense system and promote infection (Liu et al., 2014; Zhang et al., 2024).

Next, we examined how *N. benthamiana* plants responded differently when challenged with host-adapted *C. destructivum* or nonhost-adapted *C. sublineola* (**Figure 5I**). Interestingly, the two-sample t-test revealed two O-methyltransferase (OMT) proteins, which are involved in one-carbon metabolism (Lam et al., 2024), were down-regulated in plants infected with *C. destructivum.* An acetylserotonin OMT (ASMT) was one of the most down-regulated proteins (log2 FC<-8.0) and homologs are involved in the production of melatonin, which is a potent antioxidant implicated in stress responses (Back, 2021). A 3-O-methyltransferase (3-OMT) was also down-regulated and orthologs in tomato (sh3-OMT) and Arabidopsis (atOMT1) can catalyze 3’-O-methylation of flavonols, such as myricetin (Schmidt et al., 2012; Lam et al., 2024). A STRING analysis of differentially expressed proteins in *C. destructivum and C. sublineola* inoculated plants found that OMT1 belongs to a diverse group of biosynthetic proteins (**Figure 5J**), but is directly linked to flavonol synthase 1 (FLS1), a key enzyme in the flavonoid biosynthetic pathway (Owens et al., 2008). Thus, although both fungi down-regulated MATs, our data suggests the nonhost-adapted *C. sublineola*, and thus noninfectious, cannot downregulate these OMTs, potentially pointing to an NHR mode of action to prevent infection.

Multiple proteins related to jasmonic acid (JA) defense pathway were upregulated in leaves infected with *C. destructivum.* This includes an allene oxide synthase (CYP74A), which plays a crucial role in the plant’s defense against infection by initiating the biosynthesis of JA (Hughes et al., 2009; Svoboda et al., 2021). JA is a key signaling hormone that activates defense responses that includes the production of antifungal compounds, such as terpenoids and chitinases. The terpenoid biosynthetic enzyme, 5-*epi*-aristolochene synthases-like protein (EAS-like), also increased in response to *C. destructivum* and related EASs produce the antifungal terpenoid, phytoalexin capsidase (O’Maille et al., 2006). The basic chitinase (CHI-B), which is known as pathogenesis-related protein 3 (PR-3), was also upregulated in *C. destructivum* infected plants and is a well-studied antifungal enzyme that directly hydrolyzes chitin in fungal cell walls (Rawat et al., 2017). All three proteins were confidently identified by over 7 peptides, and showed significant increase upon *C. destructivum* infection (**Figure 5G**), which are consistent with previous findings. The STRING analysis (**Figure 5J**) showed that CYP74A and CHI-B belong to a defense-related group that also has pathogenesis-related protein 1 (PR1) (Mei et al., 2006). Interestingly, that group also has a SNAP25 homologous protein (SNAP33) and SYP122 syntaxin (also known as PEN1), which is a well-known fungal defense related protein (Assaad et al., 2004) found in extracellular vesicle (Rutter and Innes, 2017). SNAP33 and PEN1 are both implicated in vesicle-mediated secretion, and are vital components of a well-studied exocytic pathway for host-specific fungal disease resistance SNARE complex (Wang et al., 2016). This cluster suggests that a coordinated hormone signaling coupled exocytosis is required to reinforce pathogen restriction at the cellular interface in the plant host defense response (Yun et al., 2016).

## Discussion

In this benchmark study, we systematically investigated how an in-cell proteomics strategy could facilitate plant tissue sample preparation and showcased with comprehensive data that this on-filter in-cell digestion strategy is simple and straightforward, but also can process the “tough-to-lysis” plant tissues with exceptional depth and sensitivity. We first showed its general applicability for plant proteomics across a variety of model and crop plant species, including *Arabidopsis thaliana*, *Nicotiana benthamiana, Sorghum bicolor*, and *Zea mays*, and the ability to extract proteins from the pollen, a particularly challenging biological sample. Our OFIC approach identified ∼9,000 to 12,000 proteins from leaves and ∼7,000 to 9,000 proteins from pollen in single DIA runs. It matched or exceeded conventional lysate-based methods, without the need for mechanical disruption or detergents. The approach also bypassed the removal of highly abundant RuBisCO and related photosystem proteins, further simplifying sample preparation for plants. Lastly, OFIC can be used to study complex biological processes such the differences in plant defense responses to nonpathogenic *C. sublineola* and pathogenic *C. destructivum*.

Our OFIC approach will help democratize plant proteomics, by removing some of the complexity and technical skills required for leaf and pollen sample preparation as demonstrated in this study. The approach may also be expanded to process other plant samples such as seed, silique, flower, embryo and of course, cell culture samples. We recently have shown promising data for in-cell proteomics analysis of embryos from worms and frogs as well as data from hard-to-lysis cells such as yeast and sheath-forming bacteria (Martin et al., 2024; Keffer et al., 2025; Elsayyid et al., 2025; Sun et al., 2025b). It is worth noting that in this study, the OFIC digests were run using FAIMS and DIA LCMS, both of which are known to increase the proteome coverage in comparison to classical shotgun proteomics without FAIMS, or DDA mass spectrometry. For instance, FAIMS can reduce background noises (in particular, singly charged ions), and provide an orthogonal online gas-phase separation to conventional LCMS. It has been shown to improve identification rate (e.g., by 30-50%) by several established studies (Hebert et al., 2018; Pfammatter et al., 2018). Meanwhile, DIA compared to DDA, generally offers better quantitative accuracy, and can benefit the identification of low-abundance proteins. A very recent study by Khalil *et al*. indicated that DIA can achieve nearly 45% more proteins than DDA in the context host cell protein analysis (Khalil and Plisnier, 2025). In this regard, we speculate that the single-vessel in-cell digestion, FAIMS separation, and single-shot DIA LC-MS may all have contributed to the deep proteome analysis of plant leaf and pollen. However, because the OFIC method can generate LCMS-friendly samples in such a simple and rapid fashion with excellent yield and reproducibility, we anticipate that it can lead to even deeper proteome coverage when the OFIC digests are coupled with top-of-the-line LC-MS/MS systems.

The advent of simple in-cell proteomics in combination with one-shot FAIMS DIA LC-MS can be used to study a wide range of biological questions, including NHR and plant defense responses studied here. Although proteomics has been used extensively to study plant-microbe interactions, there are very few studies comparing non-host responses to pathogen infection. NHR is likely a combination of physical barrier defenses, pathogen-associated molecular pattern-triggered immunity, antimicrobial compounds, and effector-triggered immunity. Proteomic analysis of NHR has been limited to a few studies that compare NHR of a nonpathogenic microbe to a mock inoculation control (Li et al., 2012; Dong et al., 2015). In this study, we did a three-way comparison between the NHR of *C. sublineola*, to *C. destructivum* infection, and the mock inoculation control. Surprisingly, PEN1, which was originally identified as a key NHR component (Lipka et al., 2005), was more upregulated in *C. destructivum* infected plants than *C. sublineola* plants undergoing NHR. However, PEN1 has important roles beyond NHR in effector triggered immunity (Johansson et al., 2014) and extracellular vesicles (Rutter and Innes, 2017). Our study also discovered intriguing differences in one-carbon metabolism and implicates OMTs and involved in melatonin and flavanol production, which opens a potential future avenue of research on how they may contribute to NHR.

In conclusion, we anticipate that the simplicity and high quantitation accuracy of the “in-cell proteomics” approach presented here will allow broad researchers, in addition to plant biologists, to probe complex biological questions and identify novel regulatory pathways in an unprecedent speed and depth.

## Methods

### Plant growth, fungal treatment, and sample collection

Transgenic *Nicotiana benthamiana* plants expressing the N nucleotide-binding leucine-rich repeat (NLR) immune receptor (Liu et al., 2002) were grown in a light chamber at 150 μmol m-2 s-1 light intensity at 23 °C for 3-6 weeks until the 8-leaf stage, with a diurnal cycle of 18 hours light and 6 hours dark. Zea mays (B73) and *Sorghum bicolor* (BTX 623) seeds were grown in a 6” pot containing pro-mix BK55 growing medium for 14 days in a greenhouse or growth chamber with 16/8 h light/dark cycle at a temperature of 27 and a light intensity of 350 µmol/m^2^/s, maintaining 65% relative humidity. *Arabidopsis thaliana* (Col-0) plants were grown at 100 μmol m-2 s-1 light intensity at 20 ^O^C for 2-3 weeks, with a diurnal cycle of 16 hours light and 8 hours dark.

Cultures of *C. sublineola* and *C. destructivum* were prepared from stocks of their respective spores stored in 15% glycerol and 1X potato dextrose broth at −80 °C. Spore stocks were grown on Mathur’s media (2.8 g/L glucose, 2.2 g Mycological peptone, 1.2 g/L MgSO_4_ x 7H_2_O, 2.7 g/L KH_2_PO_4_, 30 g/L agar, pH 5.5), at 28 °C under continuous fluorescent lighting, and conidia were harvested after 7–12 d. Droplets of Conidia diluted to 2×106 conidia ml−1 were applied to *Nicotiana benthamiana* leaves using a pipette and maintained in a humid condition, stored at 28 °C. After 96 h, *N. benthamiana* leaf samples were collected and immediately stored on ice for protein extraction.

### Multiphoton microscopy and image collection

Approximately 1 mm x 1 mm leaf sections were treated with methanol or the water control for 5 minutes and then stained with cell wall dye, Calcofluor White (CW) MR for 1 minute. Multiphoton microscopy was conducted on a Zeiss LSM880 NLO microscope (Carl Zeiss, Inc, Dublin, CA) equipped with a Chameleon Discovery NX (Coherent, Inc., Santa Clara, CA) multiphoton laser tuned to 745 nm at 2% power. Fluorescence emission was collected with a band pass (BP) of 410-481 nm for CW. A z-stack with a *z*-interval size of 0.310 μm was acquired and represented in ImageJ FIJI (Schindelin et al., 2012) as a maximum intensity projection or cross-sectional views. To document chlorophyll extraction, 1 mm x 1 mm leaf sections were treated with methanol for 0, 5, 15, or 30 minutes, and then color images were acquired on an Axio Observer 7 AI (Carl Zeiss, Inc, Dublin, CA).

### Sample processing for proteomics with OFIC, SDS and TFA

For the OFIC-based sample preparation, E3filters (CDS Analytical, Oxford, PA) were used following an in-cell digestion protocol described previously (Martin et al., 2024). A detailed step-by-step protocol is provided in the **Supplement**. In brief, the leaf tissues derived from different plants were first sliced into 1-2 cm^2^ sections, and further cut into 1-2 mm^2^ small pieces, which were then transferred to E3filters followed by shaking with 400 μl of pure methanol at 1,000 rpm for 5 min. The filters were then spun at 3,000 rpm for 1 min to discard the flow through. The above shaking and spin steps were repeated 3-5 times until the green chlorophyll disappeared. The samples were then incubated with 10 mM tris (2-carboxyethyl) phosphine (TCEP) and 40 mM chloroacetamide (CAA) in 200 μl of 50 mM triethylammonium bicarbonate (TEAB) at 45 _C for 10 min, followed by a quick spin and a wash step with 200 μl of 50 mM TEAB. Unless otherwise specified, the centrifugation in this study is 3,000 rpm for 1-2 min. Afterward, the samples were transferred to clean collection tubes, and subjected to tryptic digestion in the presence of 200 μl of 50 mM TEAB and 1 μg of trypsin (Promega, WI). The samples were incubated at 37 _C for 16-18 hours with gentle shaking (300 rpm). After digestion, the filters were spun to collect flow through followed by two sequential elution, 200 μl of 0.2% formic acid (FA) in water, and 200 μl of 50% acetonitrile (ACN) in water, respectively. The elution was pooled, dried in SpeedVac, and desalted using C18-based StageTips (CDS Analytical, Oxford, PA). The dried peptides were stored under −80°C until further analysis.

For the sodium dodecyl sulfate (SDS) based preparation, the leaf pieces (1-2 mm^2^) were first lysed with 100 μl SDS buffer (4% SDS, 100 mM Tris-HCl, pH8) by vortexing at 1,500 rpm for 10 min, and then sonicated in water bath for another 10 min. The samples were then boiled with 10 mM TCEP and 40 mM CAA at 95 _C for 10 min, and cleared at 13,000 rpm for 15 min to collect the supernatant lysate. The protein lysates were processed following a standard E3 protocol reported recently (1). In brief, the lysates were mixed with 4x volume of 80% ACN to induce protein aggregation, then transferred to E3filters. Next, the filters were washed 2-3 times with 80% ACN, spun and discard the flow through. The samples were digested and desalted following the same procedure as described above. For trifluoroacetic acid (TFA) based sample preparation, the leaf pieces were incubated with 100 μl of pure TFA under room temperature for 10 min, and centrifuged at 13,000 rpm for 10 min to pellet debris. The supernatant lysates were mixed with 4x volume of pure acetone to induce protein aggregation, and transferred to E3filters. This was followed by one wash with 200 μl of acetone and another wash with 200 μl of 50 mM TEAB. The samples were then mixed with 10 mM TCEP and 40 mM CAA in 200 μl of 50 mM TEAB, and incubated at 45_C for 10 min. The E3filters were then washed once with 200 μl of 50 mM TEAB followed by the same digestion and peptide desalting procedures as described above.

For pollen sample preparation, E4tips (CDS Analytical, Oxford) was used following a similar in-cell digestion procedure reported recently (Martin et al., 2024). A detailed step-by-step protocol is provided in the **Supplement**. In brief, to collect mature pollen grains, dehiscent anthers were first gently cut off the plant and then transferred to a 1.5-ml low-binding tube, added with 500 μl of cold PBS, and then vortexed at 1,000 rpm for 5 min. The tube was centrifuged at 3,000 rpm to remove debris, and 20 μl aliquots of the resulting pollen grain suspension were transferred to E4tips that were prefilled with 200 μl of pure methanol followed by incubation at 4 _C for 15 min. A similar procedure was performed for Arabidopsis pollen sample collection, except that the pollen grains from two Arabidopsis flowers were all transferred to an E4tip. After incubation, the E4tips were spun at 4,000 rpm for 2-3 min to discard the flow through, and then subjected to reduction and alkylation by adding final concentration of 10 mM TCEP and 40 mM CAA with 200 μl of 50 mM TEAB and incubating at 45_C for 10 min. The tips were spun, washed once with 200 μl of 50 mM TEAB, and then transferred to new collection tubes. For digestion, around 1.0 μg of trypsin and 150 μl of 50 mM TEAB buffer were added to the E4tips, which were then incubated at 37 _C for 16-18 hours with gentle shaking. After digestion, about 10 μl of pure formic acid was added to the E4tips, and incubated under room temperature for 3-5 min. The tips were then centrifuged at 1,500 rpm for 10 min to allow the peptides bind to the resins, followed by a wash step with 200 μl of 0.5% acetic acid in water. The E4tips were transferred to new collection tubes, and eluted sequentially with 200 μl of 0.5% acetic acid in 60% ACN and another 200 μl of 0.5% acetic acid in 80% ACN. The elution was dried in SpeedVac and stored under −80°C until further analysis.

### LC-MS/MS

The LC-MS/MS analysis was performed using an Ultimate 3000 RSLCnano system that was coupled to an Orbitrap Eclipse mass spectrometer and FAIMS Pro Interface (Thermo Scientific). The trap column was PepMap100 C18 with a dimension of 300 μm × 2 mm, and particle size of 5 μm (Thermo Scientific). The analytical column was PepMap100 C18 (50 cm × 75 μm i.d., 3 μm; Thermo Scientific) running at a flow of 250 nl/min. A linear LC gradient was applied from 1% to 25% mobile phase B (0.1% formic acid in acetonitrile) over 125 min, followed by an increase to 32% mobile phase B over 10 min. The column was washed with 80% mobile phase B for 5 min, followed by equilibration with mobile phase A (0.1% formic acid in water) for 15 min. For MS analysis, the data-independent acquisition (DIA) mode was used. In brief, the Detector type was Orbitrap with a resolution of 60,000 for full scan; Precursor MS range (m/z) was 380–980; AGC target was Standard; Maximum injection time mode was Auto. For DIA MS/MS analysis, the Isolation mode was Quadruple; DIA Window type was Auto and Isolation Window (m/z) was 8 with an overlap of 1; activation type was HCD with fixed collision energy mode (30%); the Detector Type was Orbitrap with a resolution of 15,000; Normalized AGC target (%) was 800, and the Maximum injection time mode was Auto; the Loop Control was 2 sec. For FAIMS compensation voltages (CV) setting, a 3-CV combination (−40, −55, and −75) was applied.

### Data analysis

The MS raw data were processed using Spectronaut software (version 19.8) and a library-free DIA analysis workflow with directDIA+ and the plant proteomes obtained from UniProt Knowledgebase (Arabidopsis thaliana, taxonomy 3702, 136,324 sequences; Sorghum bicolor, taxonomy 4558, 82,122 sequences; Zea mays, taxonomy 4577, 207,460 sequences). The *N. benthamiana* protein database was curated in-house, containing 53,411 sequences from the NbDE database (Kourelis et al., 2019) combined Nicomics database (Wang et al., 2024). Detailed settings in Spectronaut included: Trypsin/P asspecific enzyme; peptide length from 7 to 52 amino acids; allowing 2 missed cleavages; toggleN-terminal M turned on; Carbamidomethyl on C as fixed modification; Oxidation on M and Acetyl at protein N-terminus as variable modifications; False discovery rates (FDRs) at PSM, peptide and protein level all set to 0.01; Quantity MS level set to MS2, and cross-run normalization turned on. Bioinformatics analyses including t-test, correlation, volcano plot and clustering analyses were performed using Perseus software (version 2.0.11.0), and Prism GraphPad (version 10.5) unless otherwise indicated. Gene Ontology analysis was conducted by using local BlastP to identify the closest Arabidopsis orthologues, which were used as the input in GeneCodis4 for cellular component GO and biological process GO (Garcia-Moreno et al., 2022). STRING protein network was built using the differentially expressed proteins between *C. destructivum* and *C. sublineola* inoculated *N.benthamiana* leaves (FDR 0.05, S0=0.1;m ≥ 2 unique peptides). The protein identifiers were converted to the accessions of *Arabidopsis thaliana* homologs obtained using BLASTP, and then imported into Cytoscape (v3.10.3) via the STRING app using the full STRING network configuration. Interactions with a confidence score ≥ 0.40 and classified as experimentally validated or curated were selected. The resulting PPI data were imported into Node degree, clustering coefficient, and betweenness centrality are computed as the network topological parameters to assess the protein connectivity. Node colors were coded according to fold changes (FC), and node sizes were weighted by p values. Functional modules were identified using the MCODE plugin and annotated based on biological function and pathway enrichment.

## Author contributions

D.A., J.L.C., and Y.Y. designed the research, interpreted data, and wrote the manuscript. D.A., M.K., and J.L.C. performed plant culturing, sample collection, imaging analysis, leaf inoculation and digestion. Y.Y. performed proteomics sample preparation and LCMS analysis. All authors analyzed the data, discussed the results and edited the manuscript.

## Acknowledgements

The authors would like to thank Mason Arnold and Jasmine Parks for the assistance of proteomics sample preparation and Tim Chaya for assistance with multiphoton microscopy. We would like to extend our special thanks to Dr. Huimin Bao from Fudan University for the assistance of image analysis and data interpretation.

## Data Availability

The MS raw data and Spectronaut result files (.sne format) associated with this study have been deposited to MassIVE server (https://massive.ucsd.edu/) via identifier MSV000097786 (password: Eclipse1).

---

**Supplementary Figure S1.**
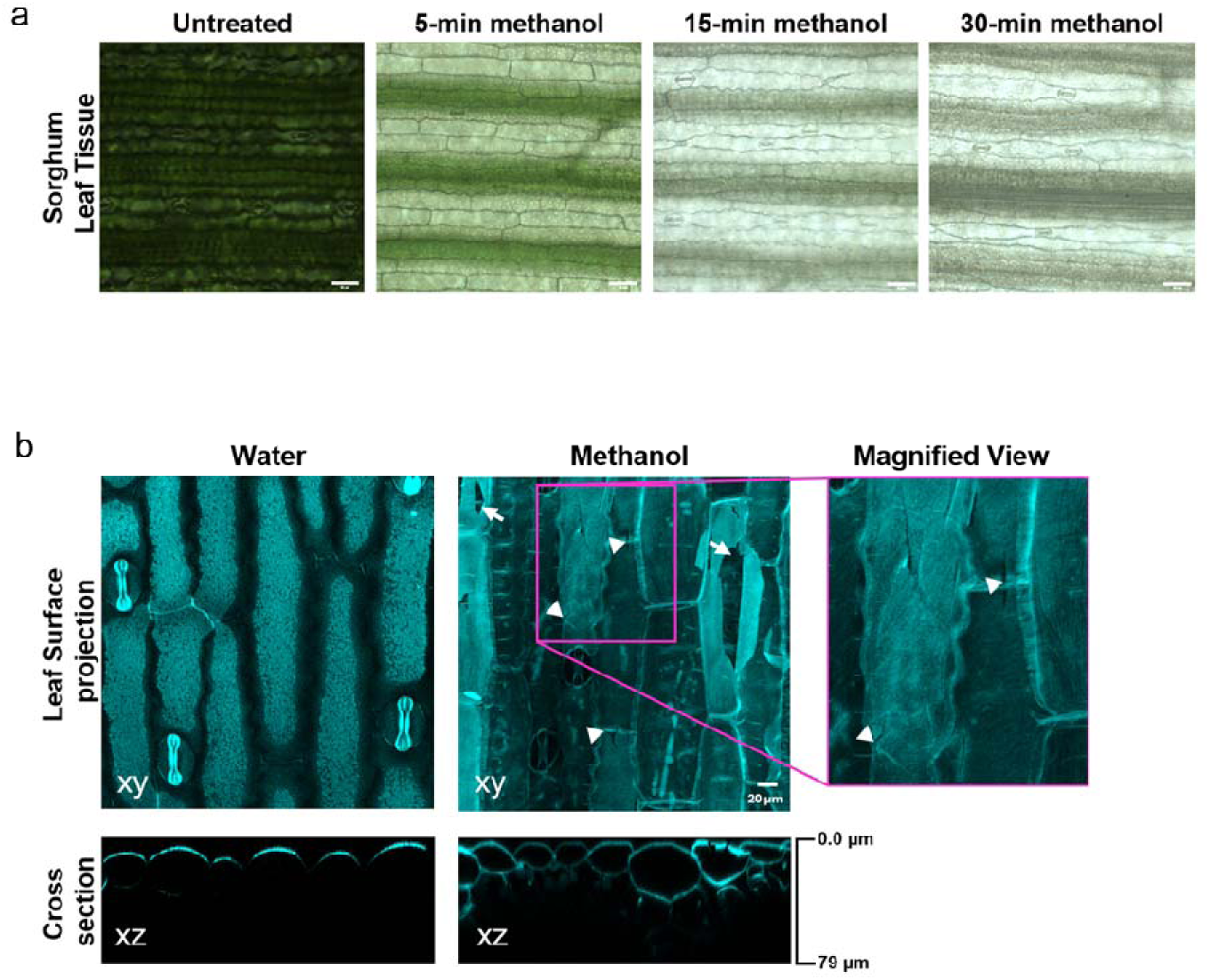
Microscopic assessment of sorghum leaf upon fixation. (**a**) Brightfield images showing the effect of 0, 5, 15, and 30 minutes of methanol treatment on leaves of *Sorghum bicolor*. Scale bar equals 50 µm. (**b**) Sorghum leaf sections treated with water or methanol, labeled with the cell wall dye Calcofluor White MR, and then imaged using multiphoton microscopy. Maximum intensity projections of the leaf surface (XY) show small cracks along the cell wall junctions (arrowheads) and larger breaks in the cell walls (arrow). A cross-sectional view (XY) comparing the staining and imaging depth between water and methanol cleared samples. Scale bar equals 20 µm.

**Supplementary Figure S2.**
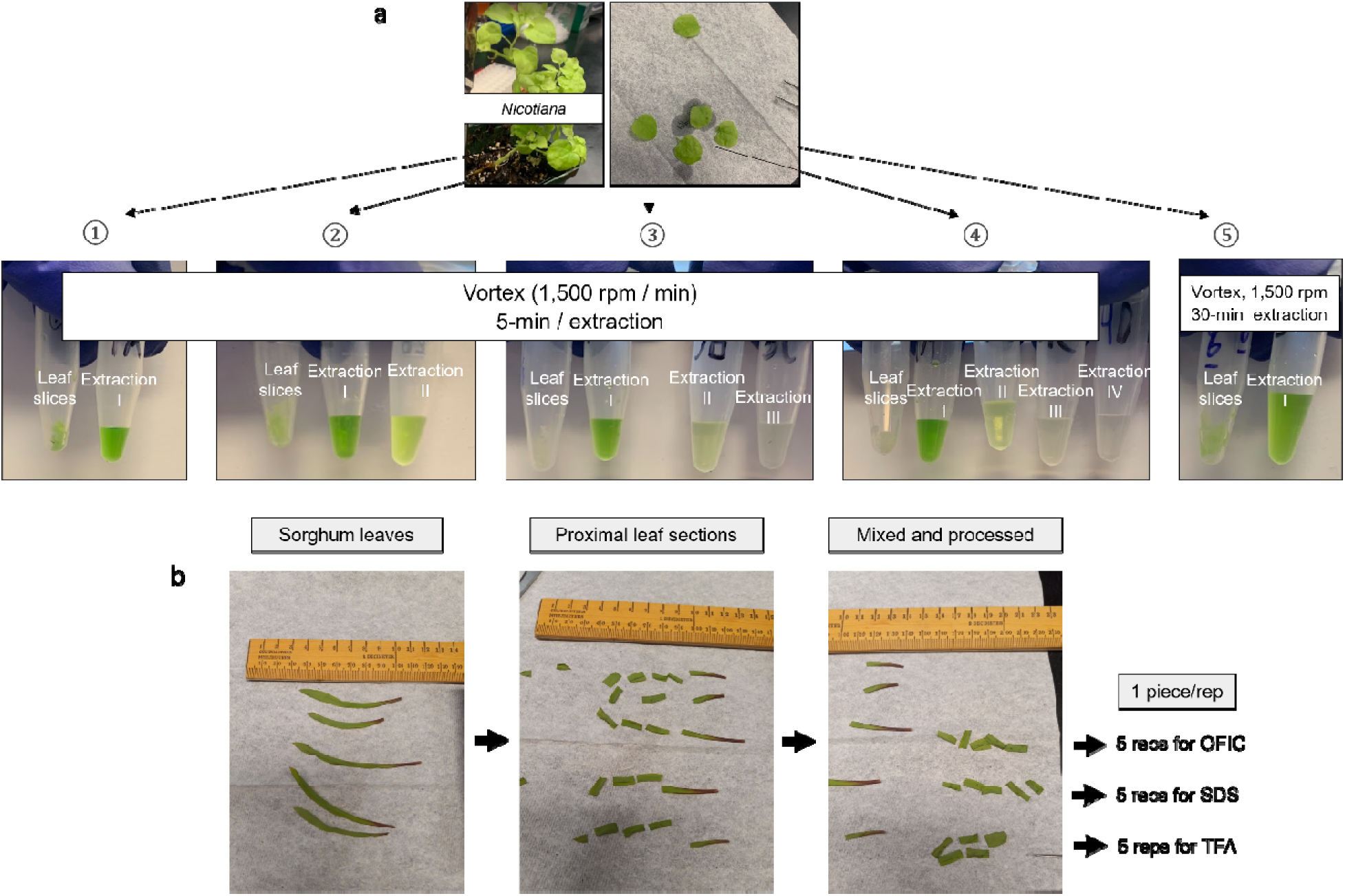
Examination of methanol fixation and leaf sample preparation. **(a)** *Nicotiana benthamiana* leaves after clearing with 1, 2, 3, or 4 exchanges (1-4) of methanol compared to a single 30-minute methanol treatment (5). **(b)** *Sorghum* leaf sample processing for proteomics analysis.

**Supplementary Figure S3.**
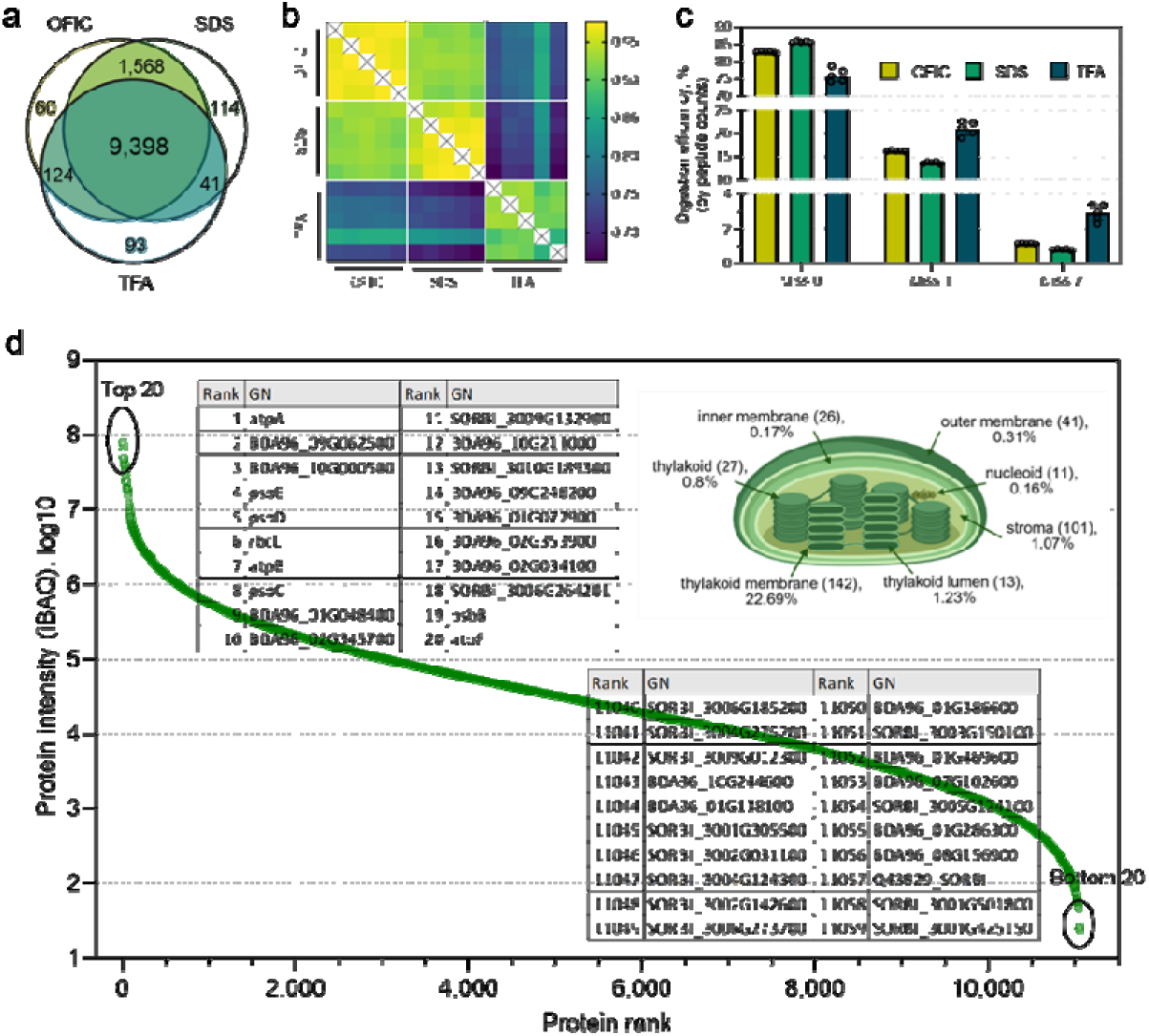
Evaluation of OFIC digestion for sorghum leaf proteomics. **(a)** Venn diagram of the overall proteins identified by each processing method. **(b)** A Pearson’s correlation analysis between replicates and treatments. **(c)** Comparison of the digestion efficiency of OFIC, SDS, and TFA methods as represented by the percentage of peptides with 0, 1, or 2 missed cleavages. (**d**) Dynamic range of sorghum leaf proteome. The 20 most and least abundant proteins (top 20 and bottom 20) are indicated on the plot. Inner panel depicts the chloroplast proteins and their relative abundance. Numbers in the brackets indicate the number of proteins of each category.

**Supplementary Figure S4.**
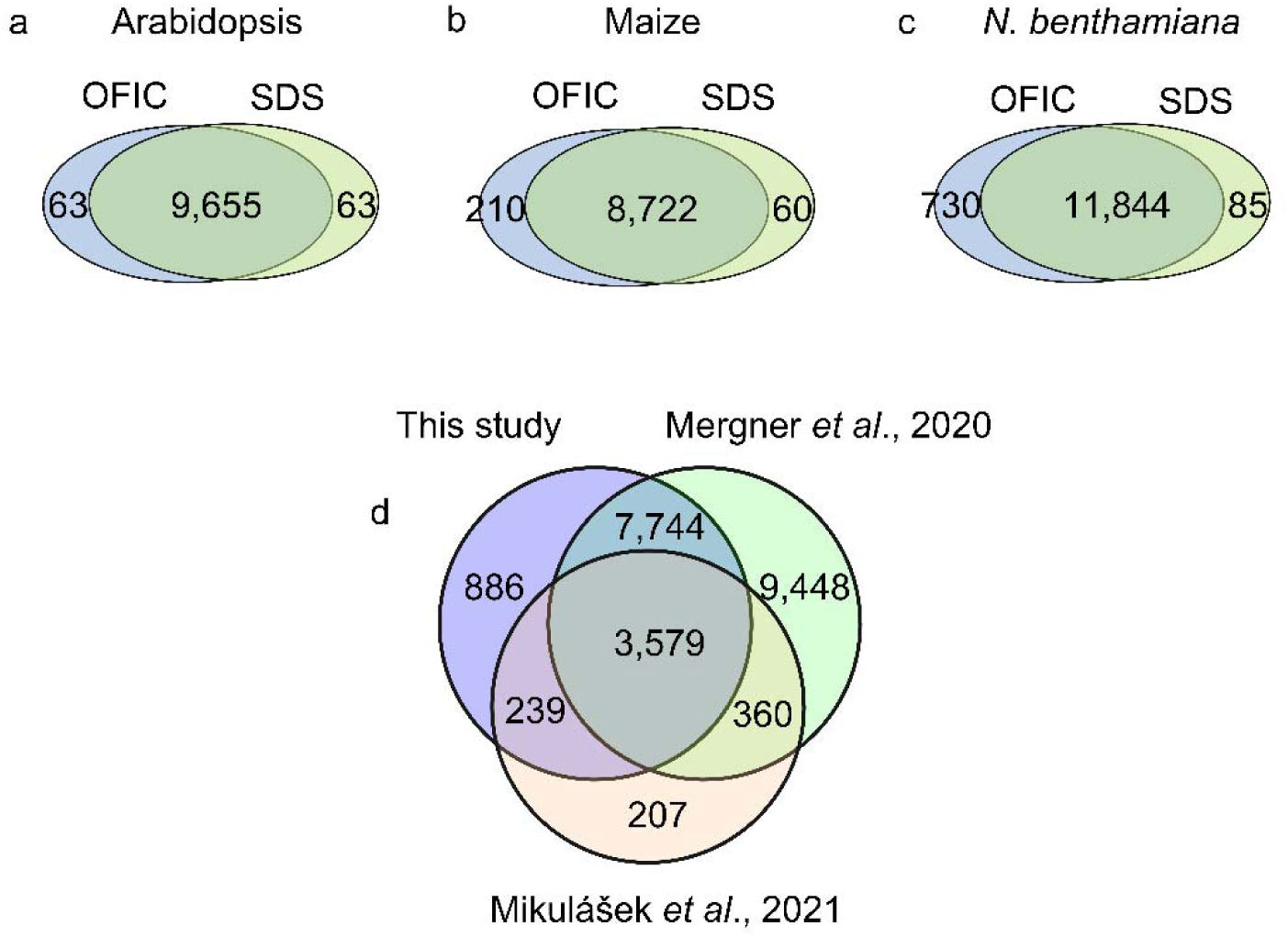
Venn diagram analysis. **(a-c)** Comparisons of protein identifications derived from OFIC and SDS digestion methods for Arabidopsis, maize, and N. benthamiana leaves, respectively. **(d)** Overlapping analysis of Arabidopsis proteins identified by this study (in-cell digestion and single-shot LCMS) with two other studies by Mergner et al., 2020 (homogenization and urea-based in-solution digestion, high pH fractionation, and multi-shot LCMS), and by Mikulášek et al. 2021 (SDS lysate and single-shot LCMS). Please be noted that the numbers used here are slightly different from the numbers reported in the main text in this study and the Mergner’s. We used all the protein accessions within each protein groups, if there are multiple members in one group, to maximize the overlaps between different studies.

**Supplementary Figure S5.**
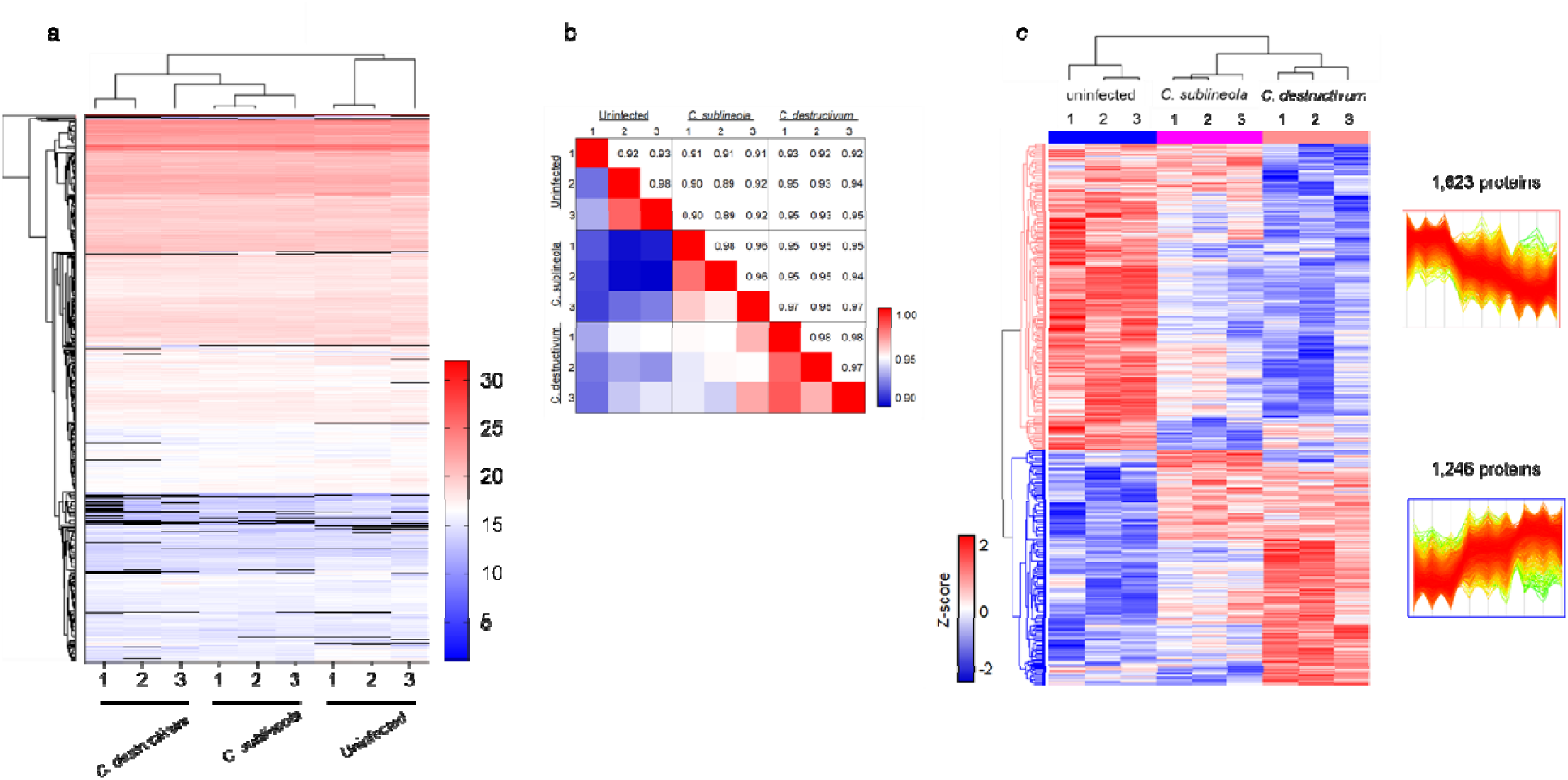
Quantitative analyses of plant leaf infection. (**a**) Unsupervised hierarchical clustering analysis of the three groups. Dark lines indicate the proteins that have missing values. (**b**) Heatmap of Pearson correlation. (**c**) Heatmap of ANOVA significant proteins among the three groups. Significance cutoff is p < 0.05. The number of proteins in the two clusters (pink and blue highlighted) are depicted.

## In-cell proteomics analysis of plant leaf using E3filter

1. Sample pretreatment and loading Collect leaves from plants; a 1-2 cm^2^ section would be sufficient for one digestion experiment. Further slice it into 1-2 mm^2^ pieces, and transfer them to E3filter (with a tweezer).
2. Methanol fixation and chlorophyl depletion Add 200-400 µl methanol, shake at 1,000 rpm for 5 min under room temperature. Centrifuge at 3,000 rpm for 1 min, discard flow through. Repeat this step (e.g.,3-6 times) until the flow through is clear with no green color.
3. Reduction and alkylation Depending on the sample size, add 100-200 µl of 50 mM triethylammonium bicarbonate (TEAB) to cover the leaf pieces, and final concentration of 10 mM Tris(2-carboxyethyl)phosphine (TCEP), incubate at 45°C for 10-15 min with gentle shaking (300-500 rpm).
4. Wash Centrifuge at 3,000 rpm for 1 min to eliminate liquid. Add 200 µl of 50 mM TEAB solution, centrifuge again, and discard flow through.
5. Digestion Transfer E3filters to clean collection tubes (here, 2-ml tube is suggested), add 150 μl of 50 mM TEAB, desired enzyme (Trypsin or Trypsin/Lys-C mix) at 1:50 ratio. Incubate at 37°C for 16-18 hours with gentle shaking. For 1-cm^2^, 1 μg of enzyme is suggested.
6. Elution After digestion, spin filters at 3,000 rpm for 1 min to collect flow through. Do two additional elutions by adding 150 μl of 0.2% formic acid in water, and 150 μl of 0.2% formic acid in 50% acetonitrile, respectively, spin and collect flow through to the same collection tube.
7. Drying and desalting Dry digests in SpeedVac; do desalting following standard StageTip protocol.

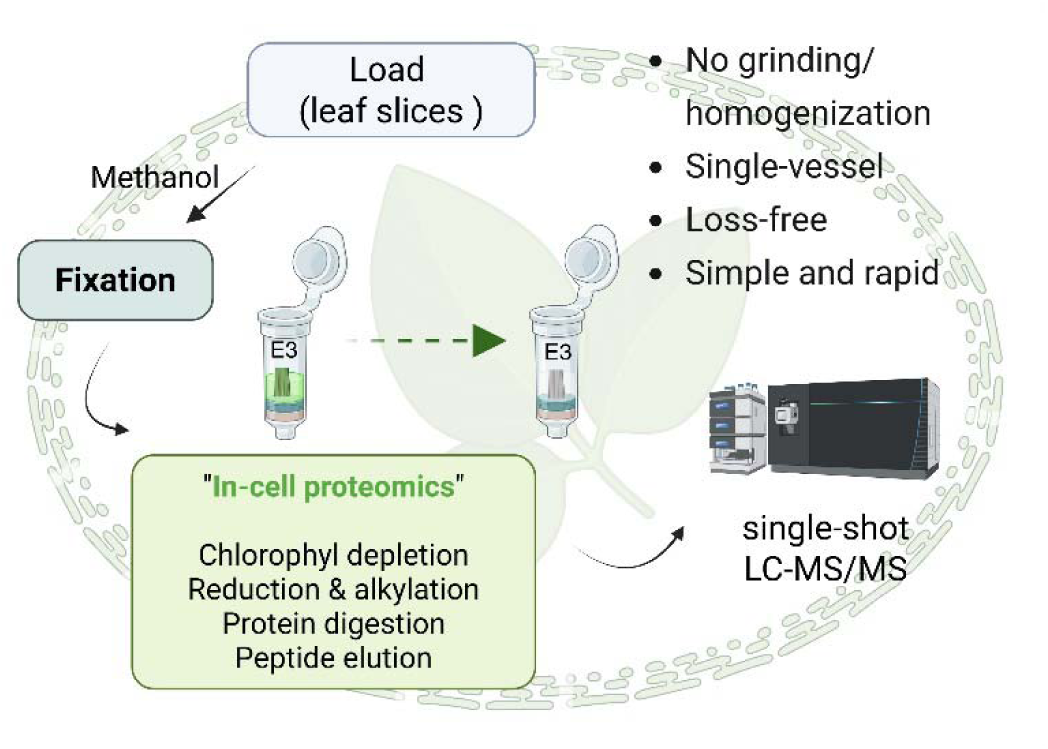

## In-cell proteomics analysis of plant pollen using E4tip

1. Sample collection and loading Collect pollen grains, aliquot 20-50 µl to E4tips that are prefilled with 150-200 µl of pure methanol.
2. Fixation Spin tips at 4,000 rpm for 2 min, discard flow through. Add 200 µl methanol to samples, and incubate at room temperature for 15 min. Centrifuge at 4,000 rpm for 2 min, discard flow through. <u>Tip</u>: the flow through may be collected here for metabolomics analysis.
3. Reduction and alkylation Add final concentration of 10 mM Tris(2-carboxyethyl)phosphine (TCEP) and 40mM chloroacetamide (CAA) in 100 µl of 50 mM triethylammonium bicarbonate (TEAB), incubate at 45°C for 10-15 min with gentle shaking (300-500 rpm).
4. Wash Centrifuge at 4,000 rpm for 1 min to eliminate liquid. Add 200 µl of 50 mM TEAB solution, centrifuge again, and discard flow through.
5. Digestion Transfer E4tips to clean collection tubes, add 150 μl 50 mM TEAB, desired enzyme (Trypsin or Trypsin/Lys-C mix) at 1:50 ratio. Incubate at 37°C for 16-18 hours with gentle shaking. <u>Tip 1</u>: please make sure no air gap between the buffer and the membrane filter. Do a pulse spin if desire (i.e., 2,000 rpm for 2 seconds). <u>Tip2</u>: No caps are required for E4tips during overnight incubation.
6. Acidification and desalting After digestion, add formic acid to final concentration of 1%, centrifuge at 1,500 rpm for 10 min. Add 200 μl 0.5% acetic acid in water, centrifuge at 4,000 rpm for 2 min, discard flow through.
7. Elution Transfer E4tips to clean collection tubes, do two sequential elution by adding 200 μl 60% acetonitrile/0.5% acetic acid in water (elution I), and 80% acetonitrile/0.5% acetic acid in water (elution II); centrifuge at 4,000 rpm for 2 min to collect eluants to the same tube. Dry samples in the SpeedVac, and store at −80°C. The peptides are now desalted and ready for LCMS analysis.

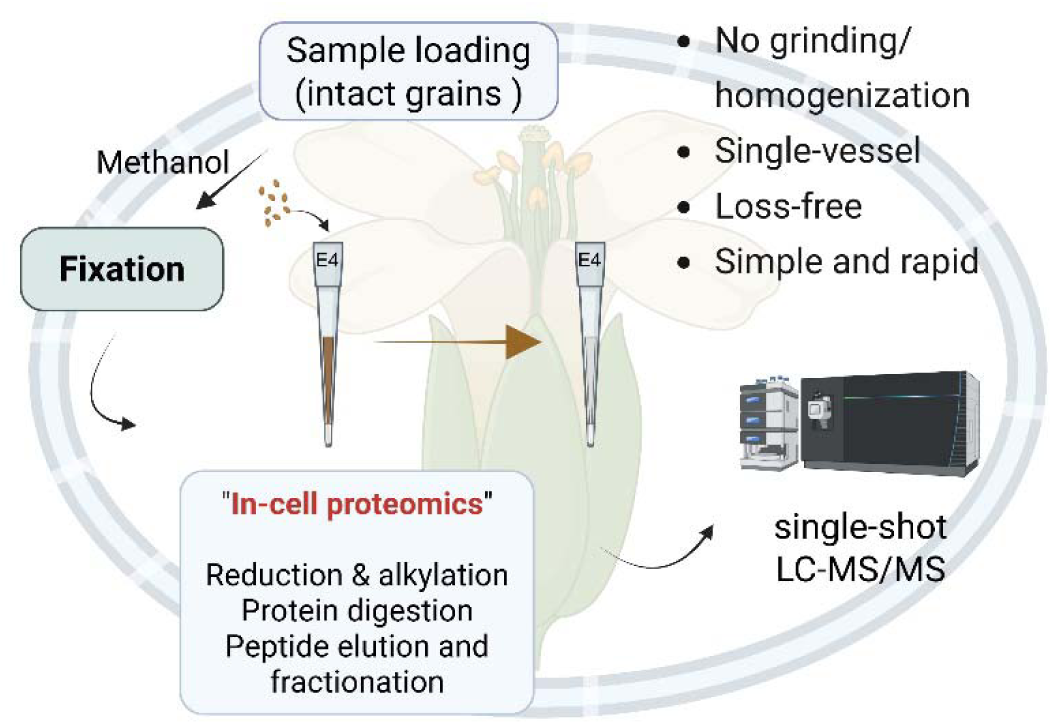

## References

Assaad, F. F., Qiu, J.-L., Youngs, H., Ehrhardt, D., Zimmerli, L., Kalde, M., Wanner, G., Peck, S. C., Edwards, H., Ramonell, K., et al. (2004). The PEN1 syntaxin defines a novel cellular compartment upon fungal attack and is required for the timely assembly of papillae. Mol. Biol. Cell 15:5118–5129.

Back, K. (2021). Melatonin metabolism, signaling and possible roles in plants. Plant J. 105:376–391.

Balotf, S., Wilson, R., Tegg, R. S., Nichols, D. S., and Wilson, C. R. (2022). Shotgun Proteomics as a Powerful Tool for the Study of the Proteomes of Plants, Their Pathogens, and Plant–Pathogen Interactions. Proteomes 10.

Bian, Y., The, M., Giansanti, P., Mergner, J., Zheng, R., Wilhelm, M., Boychenko, A., and Kuster, B. (2021). Identification of 7 000-9 000 proteins from cell lines and tissues by single-shot microflow LC-MS/MS. Anal. Chem. 93:8687–8692.

Boussardon, C., Simon, M., Carrie, C., Fuszard, M., Meyer, E. H., Budar, F., and Keech, O. (2025). The atypical proteome of mitochondria from mature pollen grains. Curr. Biol. 35:776–787.e5.

Dai, S., Chen, T., Chong, K., Xue, Y., Liu, S., and Wang, T. (2007). Proteomics Identification of Differentially Expressed Proteins Associated with Pollen Germination and Tube Growth Reveals Characteristics of Germinated Oryza sativa Pollen*S. Molecular & Cellular Proteomics 6:207–230.

Darnhofer, B., Tomin, T., Liesinger, L., Schittmayer, M., Tomazic, P. V., and Birner-Gruenberger, R. (2021). Comparative proteomics of common allergenic tree pollens of birch, alder, and hazel. Allergy 76:1743–1753.

Doellinger, J., Schneider, A., Hoeller, M., and Lasch, P. (2020). Sample Preparation by Easy Extraction and digestion (SPEED) - A universal, rapid, and detergent-free protocol for proteomics based on acid extraction. Mol. Cell. Proteomics 19:209–222.

Dong, Y., Su, Y., Yu, P., Yang, M., Zhu, S., Mei, X., He, X., Pan, M., Zhu, Y., and Li, C. (2015). Proteomic analysis of the relationship between metabolism and nonhost resistance in soybean exposed to Bipolaris maydis. PLoS One 10:e0141264.

Elsayyid, M., Tanis, J. E., and Yu, Y. (2025). Simple in-cell processing enables deep proteome analysis of low-input Caenorhabditis elegans. Anal. Chem. 97:9159–9167.

Facette, M. R., Shen, Z., Björnsdóttir, F. R., Briggs, S. P., and Smith, L. G. (2013). Parallel proteomic and phosphoproteomic analyses of successive stages of maize leaf development. Plant Cell 25:2798–2812.

Foster, A. J., Jenkinson, J. M., and Talbot, N. J. (2003). Trehalose synthesis and metabolism are required at different stages of plant infection by Magnaporthe grisea. EMBO J. 22:225–235.

Fröhlich, A., Gaupels, F., Sarioglu, H., Holzmeister, C., Spannagl, M., Durner, J., and Lindermayr, C. (2012). Looking deep inside: detection of low-abundance proteins in leaf extracts of Arabidopsis and phloem exudates of pumpkin. Plant Physiol. 159:902–914.

Gallo, M. C. R., Li, Q., Talasila, M., and Uhrig, R. G. (2023). Quantitative Time-Course Analysis of Osmotic and Salt Stress in Arabidopsis thaliana Using Short Gradient Multi-CV FAIMSpro BoxCar DIA. Molecular and Cellular Proteomics 22.

Gao, M., Liu, J., Bi, D., Zhang, Z., Cheng, F., Chen, S., and Zhang, Y. (2008). MEKK1, MKK1/MKK2 and MPK4 function together in a mitogen-activated protein kinase cascade to regulate innate immunity in plants. Cell Res. 18:1190–1198.

Garcia-Moreno, A., López-Domínguez, R., Villatoro-García, J. A., Ramirez-Mena, A., Aparicio-Puerta, E., Hackenberg, M., Pascual-Montano, A., and Carmona-Saez, P. (2022). Functional enrichment analysis of regulatory elements. Biomedicines 10:590.

Geddes-McAlister, J., and Uhrig, R. G. (2025). The plant proteome delivers from discovery to innovation. Trends Plant Sci. 30:837–845.

Gupta, R., and Kim, S. T. (2015). Depletion of RuBisCO protein using the protamine sulfate precipitation method. Methods Mol. Biol. 1295:225–233.

Gupta, R., Wang, Y., Agrawal, G. K., Rakwal, R., Jo, I. H., Bang, K. H., and Kim, S. T. (2015). Time to dig deep into the plant proteome: a hunt for low-abundance proteins. Front. Plant Sci. 6:22.

Hanson, A. D., and Roje, S. (2001). One-carbon metabolism in higher plants. Annu. Rev. Plant Physiol. Plant Mol. Biol. 52:119–137.

Hatano, A., Takami, T., and Matsumoto, M. (2023). In situ digestion of alcohol-fixed cells for quantitative proteomics. J. Biochem. 173:243–254.

Hebert, A. S., Prasad, S., Belford, M. W., Bailey, D. J., McAlister, G. C., Abbatiello, S. E., Huguet, R., Wouters, E. R., Dunyach, J.-J., Brademan, D. R., et al. (2018). Comprehensive single-shot proteomics with FAIMS on a hybrid Orbitrap mass spectrometer. Anal. Chem. 90:9529–9537.

Hughes, R. K., De Domenico, S., and Santino, A. (2009). Plant cytochrome CYP74 family: biochemical features, endocellular localisation, activation mechanism in plant defence and improvements for industrial applications. Chembiochem 10:1122–1133.

Isaacson, T., Damasceno, C. M. B., Saravanan, R. S., He, Y., Catalá, C., Saladié, M., and Rose, J. K. C. (2006). Sample extraction techniques for enhanced proteomic analysis of plant tissues. Nat. Protoc. 1:769–774.

Johansson, O. N., Fantozzi, E., Fahlberg, P., Nilsson, A. K., Buhot, N., Tör, M., and Andersson, M. X. (2014). Role of the penetration-resistance genes PEN1, PEN2 and PEN3 in the hypersensitive response and race-specific resistance in Arabidopsis thaliana. Plant J. 79:466–476.

Keffer, J. L., Zhou, N., Rushworth, D. D., Yu, Y., and Chan, C. S. (2025). Microbial magnetite oxidation via MtoAB porin-multiheme cytochrome complex in Sideroxydans lithotrophicus ES-1. Appl. Environ. Microbiol. 91:e0186524.

Khalil, S., and Plisnier, M. (2025). Definitive benchmarking of DDA and DIA for host cell protein analysis on the Orbitrap Astral in a regulatory-aligned framework. bioRxiv Advance Access published August 2, 2025, doi:10.1101/2025.07.31.667876.

Kim, Y. J., Lee, H. M., Wang, Y., Wu, J., Kim, S. G., Kang, K. Y., Park, K. H., Kim, Y. C., Choi, I. S., Agrawal, G. K., et al. (2013). Depletion of abundant plant RuBisCO protein using the protamine sulfate precipitation method. Proteomics 13:2176–2179.

Kourelis, J., Kaschani, F., Grosse-Holz, F. M., Homma, F., Kaiser, M., and van der Hoorn, R. A. L. (2019). A homology-guided, genome-based proteome for improved proteomics in the alloploid Nicotiana benthamiana. BMC Genomics 20:722.

Lam, L. P. Y., Lui, A. C. W., Bartley, L. E., Mikami, B., Umezawa, T., and Lo, C. (2024). Multifunctional 5-hydroxyconiferaldehyde O-methyltransferases (CAldOMTs) in plant metabolism. J. Exp. Bot. 75:1671–1695.

Li, H., Goodwin, P. H., Han, Q., Huang, L., and Kang, Z. (2012). Microscopy and proteomic analysis of the non-host resistance of Oryza sativa to the wheat leaf rust fungus, Puccinia triticina f. sp. tritici. Plant Cell Rep. 31:637–650.

Lipka, V., Dittgen, J., Bednarek, P., Bhat, R., Wiermer, M., Stein, M., Landtag, J., Brandt, W., Rosahl, S., Scheel, D., et al. (2005). Pre- and postinvasion defenses both contribute to nonhost resistance in Arabidopsis. Science 310:1180–1183.

Liu, Y., Schiff, M., Marathe, R., and Dinesh-Kumar, S. P. (2002). Tobacco Rar1, EDS1 and NPR1/NIM1 like genes are required for N-mediated resistance to tobacco mosaic virus. Plant J. 30:415–429.

Liu, T., Chen, J. A., Wang, W., Simon, M., Wu, F., Hu, W., Chen, J. B., and Zheng, H. (2014). A combined proteomic and transcriptomic analysis on sulfur metabolism pathways of Arabidopsis thaliana under simulated acid rain. PLoS One 9:e90120.

Ludwig, K. R., Schroll, M. M., and Hummon, A. B. (2018). Comparison of in-solution, FASP, and S-Trap based digestion methods for bottom-up proteomic studies. J. Proteome Res. 17:2480–2490.

Martin, C., Butelli, E., Petroni, K., and Tonelli, C. (2011). How can research on plants contribute to promoting human health? Plant Cell 23:1685–1699.

Martin, K. R., Le, H. T., Abdelgawad, A., Yang, C., Lu, G., Keffer, J. L., Zhang, X., Zhuang, Z., Asare-Okai, P. N., Chan, C. S., et al. (2024). Development of an efficient, effective, and economical technology for proteome analysis. *Cell Rep*. Methods 4:100796.

Mei, C., Qi, M., Sheng, G., and Yang, Y. (2006). Inducible Overexpression of a rice Allene oxide synthase gene increases the endogenous jasmonic acid level, PR gene expression, and host resistance to fungal infection. Mol. Plant. Microbe. Interact. 19:1127–1137.

Meier, F., Geyer, P. E., Virreira Winter, S., Cox, J., and Mann, M. (2018). BoxCar acquisition method enables single-shot proteomics at a depth of 10,000 proteins in 100 minutes. Nat. Methods 15:440–448.

Mergner, J., and Kuster, B. (2022). Plant proteome dynamics. Annu. Rev. Plant Biol. 73:67–92.

Mergner, J., Frejno, M., List, M., Papacek, M., Chen, X., Chaudhary, A., Samaras, P., Richter, S., Shikata, H., Messerer, M., et al. (2020). Mass-spectrometry-based draft of the Arabidopsis proteome. Nature 579:409–414.

Mikulášek, K., Konečná, H., Potěšil, D., Holánková, R., Havliš, J., and Zdráhal, Z. (2021). SP3 protocol for proteomic plant sample preparation prior LC-MS/MS. Front. Plant Sci. 12:635550.

O’Maille, P. E., Chappell, J., and Noel, J. P. (2006). Biosynthetic potential of sesquiterpene synthases: alternative products of tobacco 5-epi-aristolochene synthase. Arch. Biochem. Biophys. 448:73–82.

Owens, D. K., Alerding, A. B., Crosby, K. C., Bandara, A. B., Westwood, J. H., and Winkel, B. S. J. (2008). Functional analysis of a predicted flavonol synthase gene family in Arabidopsis. Plant Physiol. 147:1046–1061.

Pfammatter, S., Bonneil, E., McManus, F. P., Prasad, S., Bailey, D. J., Belford, M., Dunyach, J.-J., and Thibault, P. (2018). A novel differential ion mobility device expands the depth of proteome coverage and the sensitivity of multiplex proteomic measurements. Mol. Cell. Proteomics 17:2051–2067.

Raskin, I., Ribnicky, D. M., Komarnytsky, S., Ilic, N., Poulev, A., Borisjuk, N., Brinker, A., Moreno, D. A., Ripoll, C., Yakoby, N., et al. (2002). Plants and human health in the twenty-first century. Trends Biotechnol. 20:522–531.

Rawat, S., Ali, S., Mittra, B., and Grover, A. (2017). Expression analysis of chitinase upon challenge inoculation to Alternaria wounding and defense inducers in Brassica juncea. Biotechnol. Rep. (Amst.) 13:72–79.

Rejón, J. D., Delalande, F., Schaeffer-Reiss, C., de Alché, J. D., Rodríguez-García, M. I., Van Dorsselaer, A., and Castro, A. J. (2016). The pollen coat proteome: At the cutting edge of plant reproduction. Proteomes 4:5.

Rutter, B. D., and Innes, R. W. (2017). Extracellular Vesicles Isolated from the Leaf Apoplast Carry Stress-Response Proteins. Plant physiology 173:728–741.

Sage, R. F. (2002). Variation in the k(cat) of Rubisco in C(3) and C(4) plants and some implications for photosynthetic performance at high and low temperature. J. Exp. Bot. 53:609–620.

Schindelin, J., Arganda-Carreras, I., Frise, E., Kaynig, V., Longair, M., Pietzsch, T., Preibisch, S., Rueden, C., Saalfeld, S., Schmid, B., et al. (2012). Fiji: an open-source platform for biological-image analysis. Nat. Methods 9:676–682.

Schmidt, A., Li, C., Jones, A. D., and Pichersky, E. (2012). Characterization of a flavonol 3-O-methyltransferase in the trichomes of the wild tomato species Solanum habrochaites. Planta 236:839–849.

Shen, J.-J., Chen, Q.-S., Li, Z.-F., Zheng, Q.-X., Xu, Y.-L., Zhou, H.-N., Mao, H.-Y., Shen, Q., and Liu, P.-P. (2022). Proteomic and metabolomic analysis of Nicotiana benthamiana under dark stress. FEBS Open Bio 12:231–249.

Song, G., Hsu, P. Y., and Walley, J. W. (2018). Assessment and refinement of sample preparation methods for deep and quantitative plant proteome profiling. Proteomics 18:e1800220.

Sun, J., Xu, X., and Wei, S. (2025a). -cell Proteomics Enables High-Resolution Spatial and Temporal Mapping of Early Xenopus tropicalis Embryos.

Sun, J., Xu, X., Wei, S., and Yu, Y. (2025b). In-cell proteomics enables high-resolution spatial and temporal mapping of early Xenopus tropicalis embryos. bioRxivorg Advance Access published May 28, 2025, doi:10.1101/2025.05.23.655823.

Svoboda, T., Thon, M. R., and Strauss, J. (2021). The role of plant hormones in the interaction of Colletotrichum species with their host plants. Int. J. Mol. Sci. 22:12454.

Tola, A. J., and Missihoun, T. D. (2023). Ammonium sulfate-based prefractionation improved proteome coverage and detection of carbonylated proteins in Arabidopsis thaliana leaf extract. Planta 257:62.

Wang, W.-M., Liu, P.-Q., Xu, Y.-J., and Xiao, S. (2016). Protein trafficking during plant innate immunity. J. Integr. Plant Biol. 58:284–298.

Wang, W.-Q., Jensen, O. N., Møller, I. M., Hebelstrup, K. H., and Rogowska-Wrzesinska, A. (2018). Evaluation of sample preparation methods for mass spectrometry-based proteomic analysis of barley leaves. Plant Methods 14:72.

Wang, L., Lau, Y.-L., Fan, L., Bosch, M., and Doughty, J. (2023a). Pollen Coat Proteomes of Arabidopsis thaliana, Arabidopsis lyrata, and Brassica oleracea Reveal Remarkable Diversity of Small Cysteine-Rich Proteins at the Pollen-Stigma Interface. Biomolecules 13:157.

Wang, L., Patena, W., Van Baalen, K. A., Xie, Y., Singer, E. R., Gavrilenko, S., Warren-Williams, M., Han, L., Harrigan, H. R., Hartz, L. D., et al. (2023b). A chloroplast protein atlas reveals punctate structures and spatial organization of biosynthetic pathways. Cell 186:3499–3518.e14.

Wang, J., Zhang, Q., Tung, J., Zhang, X., Liu, D., Deng, Y., Tian, Z., Chen, H., Wang, T., Yin, W., et al. (2024). High-quality assembled and annotated genomes of Nicotiana tabacum and Nicotiana benthamiana reveal chromosome evolution and changes in defense arsenals. Mol. Plant 17:423–437.

Wu, X., Xiong, E., Wang, W., Scali, M., and Cresti, M. (2014). Universal sample preparation method integrating trichloroacetic acid/acetone precipitation with phenol extraction for crop proteomic analysis. Nat. Protoc. 9:362–374.

Xu, X., Yu, Y., Zheng, T., Clark, F., Ross, J., Tavares, A. L. P., Wei, S., and Sun, J. (2025). Beyond the axoneme: Whole-cell proteomics identifies novel regulators of ciliogenesis in Xenopus epidermal multiciliated cells. bioRxivorg Advance Access published May 21, 2025, doi:10.1101/2025.05.20.655211.

Yun, H. S., Kang, B. G., and Kwon, C. (2016). Arabidopsis immune secretory pathways to powdery mildew fungi. Plant Signal. Behav. 11:e1226456.

Zhang, H., Liu, P., Guo, T., Zhao, H., Bensaddek, D., Aebersold, R., and Xiong, L. (2019). Arabidopsis proteome and the mass spectral assay library. bioRxiv Advance Access published June 11, 2019, doi:10.1101/665547.

Zhang, C., Li, H., Yin, J., Han, Z., Liu, X., and Chen, Y. (2024). Pan-genome wide identification and analysis of the SAMS gene family in sunflowers (Helianthus annuus L.) revealed their intraspecies diversity and potential roles in abiotic stress tolerance. Front. Plant Sci. 15:1499024.

Zheng, K., Lyu, J. C., Thomas, E. L., Schuster, M., Sanguankiattichai, N., Ninck, S., Kaschani, F., Kaiser, M., and van der Hoorn, R. A. L. (2024). The proteome of Nicotiana benthamiana is shaped by extensive protein processing. New Phytol. 243:1034–1049.

